# Differential Left and Right Carotid Artery Blood Flow and Altered Hippocampal Mitochondrial Function After Transverse Aortic Constriction in Aging Rats

**DOI:** 10.1101/2025.01.15.633217

**Authors:** Gabriel S. Pena, Yuan Liu, Maria Canellas Da Silva, Sushant Ranadive, J. Carson Smith, Sarah Kuzmiak-Glancy

## Abstract

**Background:** The hippocampus is a key brain structure that has been implicated in vascular dementia etiology and is highly sensitive to changes in cerebral blood flow. Brain hypoperfusion in cardiovascular disease may facilitate neurodegeneration in the hippocampus by limiting substrate transport and metabolism. While most animal studies have relied on artery occlusion to lower brain blood flow, brain hypoperfusion can also stem from mechanical damage resulting from high blood flow velocity and pulsatility. This study assessed, within the same rodent, whether high and low cerebral blood flow differentially affected hippocampal glucose transport protein expression, mitochondrial fuel oxidation, and expression of mitochondrial quality control proteins.

**Methods and Results:** 4-week-old male and female Sprague-Dawley rats underwent transverse aortic constriction (TAC, n=13) or control (SHAM, n=18) surgeries. Bilateral carotid artery diameter and blood flow were measured 20-, 30-, and 40 weeks post-surgery. Right and left hippocampal mitochondrial respiration and expression of glucose transporters and mitochondrial quality control proteins were measured 40 weeks post-surgery. Right carotid blood flow velocity and pulsatility were highest in the right and lowest in the left carotid of TAC animals (p<0.05). Complex I (p=0.057), Complex I&II (p<0.05), and Complex II uncoupled (p<0.05) respiration rates were lower in the right hippocampus of TAC when compared to the left, and markers of mitochondrial fusion were upregulated in TAC compared to SHAM (p<0.05).

**Conclusions:** While both limited and pulsatile blood flow alter mitochondrial fusion markers, only pulsatile flow impairs mitochondrial respiration, suggesting turbulent hemodynamics may drive metabolic dysfunction in vascular dementia.

**Clinical Perspective:** *What is new?:*

- High blood velocity and pulsatility impairs hippocampal mitochondrial respiration while reduced blood flow does not.
- Hippocampal protein expression of mitochondrial fusion and unfolded protein response markers is upregulated in response to altered carotid blood flow, but protein expression of endothelial and neuronal glucose transporters is unaffected.
- Hippocampal mitochondrial respiration and protein expression of mitochondrial dynamics markers differs between male and females.

*What are the clinical implications?:*

- Brain hypoperfusion stemming from high blood velocity and pulsatility negatively impacts glucose oxidation by brain mitochondria to a greater degree than hypoperfusion stemming from decreased brain blood flow, and optimal treatment of vascular dementia may be best informed by vascular hemodynamic phenotype rather than cortical perfusion alone.
- Alterations in mitochondrial structure and function may precede changes in glucose transport during brain hypoperfusion, demonstrating the potential importance of treating mitochondrial impairments in vascular dementia.
- **Nonstandard Abbreviations and Acronyms**

## Introduction

Cardiovascular and neurodegenerative diseases represent two of the most common age-related chronic diseases among older adults^1–3^ that cumulatively account for 36% of the disease burden in the older adult population^4^. Cardiovascular health and brain health are closely coupled^5^, and cardiovascular disease can increase the risk of developing neurodegenerative diseases^6^ and exacerbate the rate of progression and clinical manifestations of dementia^7^. In fact, vascular pathologies can independently give rise to a spectrum of deficits in brain function and structure clinically known as vascular cognitive impairment (VcI) and vascular dementia (VaD)^8^. Nevertheless, the cellular mechanisms by which neurodegeneration can be initiated, influenced, or exacerbated by cardiovascular disease remain poorly understood^9, 10^.

Chronic brain hypoperfusion is widely seen as a key link between cardiovascular disease and neurodegeneration^11^. Human studies show impaired brain blood flow^12^ and brain perfusion^13, 14^ may lower oxygen and glucose delivery to neural mitochondrial and facilitate severe energetic deficits^15^. Although brain hypoperfusion is seldomly operationally defined, it can be a consequence of two distinct hemodynamic phenotypes^16^. The first rises from conditions that lower circulating blood volume (heart failure) or blood pressure (orthostatic hypotension) and lead to lower blood volume and/or insufficient perfusion pressures^16, 17^. The second stems from conditions that lower arterial compliance and increase blood flow velocity and pulsatility (e.g., hypertension, carotid stenosis, etc.), that ultimately lead to chronic mechanical strain and end-organ damage^18^. Both phenotypes can result in lower brain perfusion and brain glucose uptake^19, 20^. Lower perfusion and glucose uptake could negatively impact neural mitochondrial ATP production^15, 21^. Thus, the disruption of glucose transport and altered mitochondrial ATP production in chronic brain hypoperfusion may be an underlying mechanism linking cardiovascular disease to neurodegeneration.

A major limitation in studying brain hypoperfusion in animal models of cardiovascular disease and neurodegeneration is the reliance on unilateral or bilateral artery occlusion to obstruct blood flow^22, 23^. These models fail to recognize that brain hypoperfusion can also arise from high blood flow velocity and pulsatility^18^. Although both mechanisms of brain hypoperfusion have been independently reported to negatively impact neuronal mitochondria^24, 25^, the magnitude of tissue hypoperfusion and the severity of cerebrovascular dysfunction that follow are not equivalent between the two^26^. Therefore, while it is likely that these contrasting hypoperfusion mechanisms can exert distinct effects on the cellular metabolic processes linking cardiovascular disease and neurodegeneration (e.g., neuronal glucose transport and mitochondrial metabolism), it remains a gap of knowledge that remains unaddressed in the literature.

To investigate the metabolic consequences of the two hemodynamic phenotypes of brain hypoperfusion, we utilized transverse aortic constriction (TAC) in male and female rats given it can generate the two hemodynamic phenotypes within the same animal. TAC involves the partial ligation of the aorta between the right and left common carotid arteries, inducing high pulsatile blood flow through the right carotid while limiting blood flow through the left carotid, a response that is paralleled in the cerebral hemispheres^23, 27^. As such, there were two primary aims for this study. The first was to characterize the long-term hemodynamic response to TAC (20-, 30-, and 40-weeks post-surgery) in the left and right carotid arteries of male and female rats. The second was to investigate whether mitochondrial respiration, content, and protein expression of glucose transporters and mitochondrial quality control markers differed in the left and right hippocampus in TAC rats, and if these differed from SHAM-operated controls.

## Methods

This study was approved by the Institutional Animal Care and Use Committee (IACUC) at the University of Maryland College Park. Male and female Sprague-Dawley rats underwent either transverse aortic constriction (TAC) surgery to mimic chronic hypertension and induced heart failure with preserved ejection fraction or a SHAM surgery, that was identical to TAC, barring the constriction of the aorta. Following surgical procedures and wound healing, animals were paired-housed, fed water and food ad libitum and aged for 40 weeks.

### Transverse Aortic Constriction Surgery

Male and female Sprague-Dawley rats (4 weeks old) underwent either the TAC or SHAM surgical procedures as described previously^28^. Briefly, each animal was weighed and anesthetized via inhalation of isoflurane [∼2% isoflurane supplemented with 100% oxygen (400-500 mL/min)]. Once unresponsive, a single dose of buprenorphine (0.05-0.1 mg/kg) was injected subcutaneously, and the surgical area was clipped clean and sterilized. A horizontal skin incision was made, at the suprasternal notch region, a longitudinal ∼1 cm cut was made in the sternum, and the left and right carotids were visualized. In both procedures, a 4-0 silk suture attached to a blunted needle was passed around the aortic arch between the origin of the right innominate and left common carotid arteries. A bent, blunted 20-gauge needle was then placed next to the aortic arch, the suture was tightened around the needle, and the needle was then promptly removed. The sternum was closed with a single silk suture and the skin was sutured shut with 5-0 monofilament suture and treated with betadine. SHAM animals received the same procedure as TAC animals but did not have the suture tied around the aortic arch.

### Cardiac and Carotid Doppler Ultrasound

Carotid blood flow and artery diameter were measured with a Vevo3100 (FUJIFILM visualsonics) animal Doppler ultrasound imaging system at 20-, 30-, and 40-weeks post-surgery. All animals were placed under anesthesia in an induction chamber with 2-5% isoflurane supplemented with 100% oxygen (400-500 mL/min). Once animals became unresponsive, the fur on the chest was removed, ophthalmic gel was applied to both eyes, and animals were placed on an imaging platform and administered isoflurane via nose cone (∼2% isoflurane supplemented with 100% oxygen (400-500 mL/min)). Ultrasound imaging of the aortic arch and cardiac tissue were used to confirm aortic arch binding and obtain cardiac parameters such as heart rate, cardiac output, ejection fraction, and stroke volume under anesthesia. Carotid artery blood flow was determined with the use of B-mode & PW Doppler Imaging of the bilateral carotid arteries (**Fig. 1A**). Ultrasound files (PW Doppler and B-mode) were analyzed in triplicates with the Vevolab software (FUJIFILM visualsonics) and carotid arterial blood flow velocity (mean and peak velocities), blood flow pulsatility (pulsatility index), and artery diameter during systole and diastole were determined.

**Figure 1.**
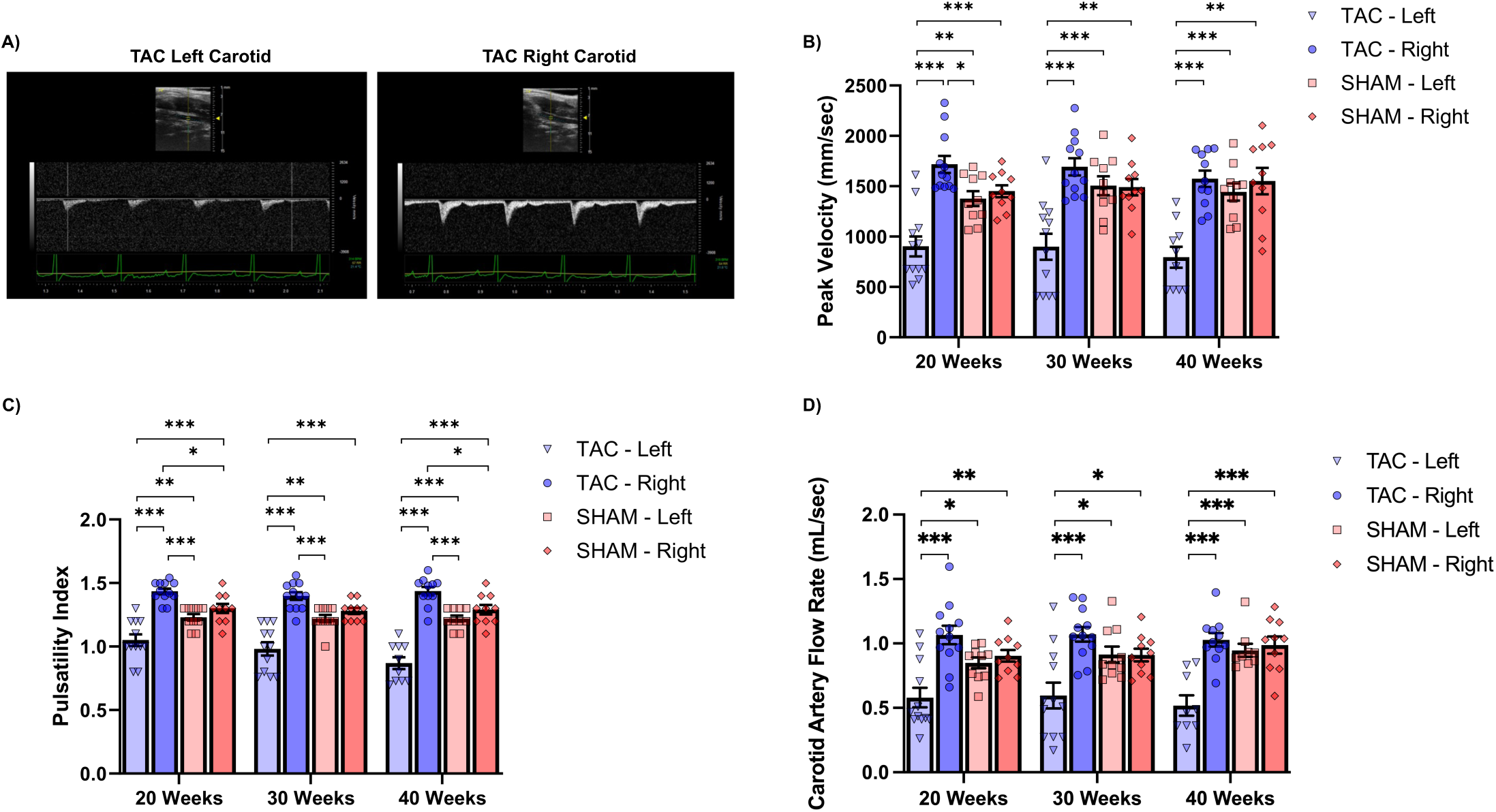
TAC carotid hemodynamic phenotype is characterized by greater right and lower left carotid blood flow that does not worsen with time: **A)** Doppler ultrasound waveforms of the left and right carotid arteries of TAC Animals. **B)** Peak carotid blood flow velocity in the right and left carotids of TAC and SHAM animals 20-, 30-, and 40 weeks post-surgery. **C)** Blood flow pulsatility in the right and left carotid of TAC and SHAM animals 20-, 30-, and 40 weeks post-surgery. **D)** Calculated right and left carotid flow rate in TAC and SHAM animals 20-, 30-, and 40 weeks post-surgery. TAC = transverse aortic constriction, SHAM = sham control, mm/sec- millimeter per second, µl/sec- microliter per second, * = p<0.05, ** = p<0.01, *** = p<0.001. Data presented as mean + SEM.

### Euthanasia and Tissue Isolation

Rats were first anesthetized (∼2% isoflurane supplemented with 100% oxygen (400-500 mL/min)). Once unresponsive, the heart was removed via bilateral thoracotomy, the head was removed, and the brain was dissected out. Whole brain samples were placed in ice-cold 1xPBS, and the bilateral hippocampi were isolated and weighed. The hippocampus was selected as a structure of interest because it is a subcortical structure particularly sensitive to neurodegenerative diseases^29, 30^, and is one that is heavily targeted in the animal literature of brain hypoperfusion^23, 25, 26^. Following bilateral hippocampal isolation, ∼11mg of the right and left hippocampus were homogenized in 4.4 mL of ice-cold respiration media (50mM MOPS, 20mM glucose, 1mM EGTA, 100mM KCl, 10mM MgCl_2_, 0.2% BSA) using separate ice-cold Dounce homogenizers. The remaining hippocampal samples were placed in separate conical tubes with ice-cold 1xRIPA lysing buffer containing protease and phosphatase inhibitors (ThermoFisher Scientific, Waltham, MA) and homogenized using a hand-held motorized microtube homogenizer (VWR, Radnor, PA). Samples were spun at 15xG for 15 minutes to obtain purified protein from each sample, which were then stored at −80°C for further analysis.

### Hippocampal Respiration

Hippocampal mitochondrial oxygen consumption was obtained via liquid-phase respiration in a temperature-controlled respiration chamber fitted with a Clark-type electrode (Oxytherm^+^, Hansatech Instruments, Norfolk, UK). 1 mL of hippocampal homogenate was placed into the respiration chamber with the temperature preset at 37°C with constant stirring at 60 rpm, and a previously published substrate-uncoupler-inhibitor-titration (SUIT) respiration protocol was used to assess mitochondrially-fueled metabolism^31^. While continually monitoring oxygen consumption, non-ADP-phosphorylating leak respiration was induced by adding the CI-linked substrates pyruvate (5 mM), malate (0.5 mM) and glutamate (10 mM). Then ADP (2.5 mM) was added to measure maximal CI-linked, ADP-phosphorylating oxygen consumption, and then succinate (10 mM) was added, to assess the combined CI + CII phosphorylating oxygen consumption rate. Maximal uncoupled respiration was obtained with the stepwise titration of protonophore FCCP (0.5 μM additions separated by 60 seconds), which dissipates the mitochondrial protonmotive force, and allows for an estimation of maximal ETS electron transportation capacity. Rotenone (0.5 μM) was then used to inhibit CI and obtain CII-linked uncoupled respiration. Lastly, CII and CIII were inhibited with malonate (2mM) and antimycin A (2.5mM) to assess residual non-mitochondrial oxygen consumption (ROX).

### Hippocampal Mitochondrial Content

Citrate synthase activity was assessed as an estimate of mitochondrial content and was performed following previously published protocols^32^. Hippocampal homogenates were added to a cuvette containing, 100 mM Tris HCl, 0.1 mmol 5,5-dithiobis-(2-nitrobenzoic acid) (DTNB), 0.25mM acetyl-CoA, and 0.1% Triton, and background was read. Substrate dependent activity was initiated by addition of 0.5 mM oxaloacetate into a final volume of 1.0 ml and absorbance was followed at 412nm for 180 seconds. Activity was calculated using a millimolar extinction coefficient of 13.6 for the mercaptide ion.

### Western Blotting

Right and left hippocampal protein concentrations were first determined via Pierce BCA protein assay (ThermoFisher Scientific, Waltham, MA). Equal amounts of protein (30 μg/lane) were loaded and separated by SDS-PAGE using Mini-PROTEAN stain-free TGX precast gels (2% SDS, 25% glycerol, 0.01% bromophenol blue) (Bio-Rad, Hercules, CA), and transferred to PVDF membranes (Bio-Rad, Hercules, CA). Membranes were then blocked in 5-6% nonfat dry milk in TBST for one hour and then incubated at 4°C overnight in 3% nonfat dry milk in TBST with primary antibodies for target proteins. The primary antibodies for this study included: GLUT-1 (Cell Signaling #12939, 1:900), GLUT-3 (Invitrogen #MA532697, 1:900), DRP1 (Cell Signaling #8570, 1:1000), FIS1 (Millipore Sigma #HPA017430, 1:900), MFN2 (Cell Signaling #83667, 1:1500), OPA1 (Cell Signaling #80471, 1:1000), and HSP60 (Cell Signaling #12165, 1:1600). An HRP-conjugated secondary antibody was used (Cell Signaling #7074), and the SuperSignal West Pico Chemiluminescent Substrate detection kit was used to initiate the immunoreactive protein reaction (ThermoFisher Scientific, Waltham, MA). Protein bands were detected and imaged with the Bio-Rad ChemiDoc system and software (Bio-Rad, Hercules, CA), and ImageJ software was used to quantify bands using densitometry. For all western blot experiments, total protein was used as the normalizing control and was obtained using stain-free imaging technology (Bio-Rad, Hercules, CA).

### Statistical Analysis

To test any main effects of Condition (TAC v. SHAM), Hemisphere (right v. left), or Condition by Hemisphere interactions on carotid blood flow, pulsatility index, and artery diameter, a series of Independently run 2×2 ANOVA were used and significant Condition by Hemisphere interactions were followed up with a post-hoc analysis employing Tukey’s HSD correction for multiple comparisons. Additionally, independently run one-way ANOVA models were used to determine whether there were any main effects of Time (20-, 30-, and 40 weeks) on carotid blood flow, pulsatility index, and artery diameter measures. Separate 2×2 ANOVA models were also used to test whether there were any main effects of Condition (TAC v. SHAM), Hemisphere (right v. left), or Condition by Hemisphere interactions on hippocampal respiration, mitochondrial content, and protein expression of markers associated with mitochondrial quality control. Because previous literature has found hemispheric differences in brain perfusion within TAC, and because we also hypothesized these hemispheric differences would also be present for mitochondrial respiration, and protein expression of target markers, all 2×2 ANOVA models for mitochondrial outcomes were followed up with planned comparisons to specifically test whether there were any hemispheric differences within TAC. Pearson product-moment correlations were used to determine if carotid artery blood flow velocities and pulsatility 40 weeks post-surgery were related to mitochondrial respiration. Lastly, separately run, independent samples t-test, or Mann-Whitney U test, were used as exploratory analyses to address any possible sex differences on right/left carotid blood flow, pulsatility index, artery diameter measures, hippocampal respiration, mitochondrial content, and protein expression within either the TAC or SHAM conditions. All analyses were conducted using the statistical software JASP (version 0.17.3), and statistical significance was set at p < 0.05. Data that support the findings of this study are available from the corresponding author upon reasonable request.

### Excluded and Missing Data

Thirty-one animals were used in the investigations (7 TAC male, 6 TAC female, 7 SHAM male, and 10 SHAM female). However, for the assessment of carotid hemodynamics, a total of 1 TAC (1 female) and 8 SHAM (4 male, 4 female) animals were excluded from the analysis due to poor ultrasound quality or missing data. Further, Doppler ultrasound time series revealed 3 TAC animals had left carotid blood flow obstructed to such degree that blood flow wave forms could not be analyzed (e.g., flat wave forms) despite good image contrast and measurable flows on the right carotid (**Supplemental Figure 1A**). Instead of assigning a blood flow value of “0” for these animals, the lowest measured left carotid values within the TAC group were assigned as proxy left carotid values to provide a realistic, conservative flow value. Lastly, 2 male TAC animals with complete data at 20-, and 30-weeks but incomplete data at 40 weeks (due to poor ultrasound image resolution) were included in the analyses for which the respective data was available (e.g., 20- and 30 weeks 2×2 ANOVA) but excluded from the timepoints for which no data was available (e.g., 40 weeks 2×2 ANOVA). As such, for the carotid hemodynamic assessments only, a total of 12 TAC (7 male, 5 female) and 10 SHAM (4 male, 6 female) with a complete set of data for most time points were utilized. For all other study outcomes, data was obtained from 30 animals - as respiration data for 1 TAC (female) and western blot data for 1 SHAM (male) could not be obtained due to an unstable oxygen trace and protein degradation, respectively. Animal characteristics for the full (n=31) cohort can be found in Table 1. Overall, no significant differences between TAC and SHAM were observed in any of the characteristics, including hippocampal mitochondrial content, heart mass, septal width, cardiac output, ejection fraction, and stroke volume (**Table 1**).

**Table 1:** Animal Characteristics.

|  | TAC (n=13)<br>M (SEM) | SHAM (n=18)<br>M (SEM) |
| --- | --- | --- |
| Sex |  |  |
| Male | 7 | 8 |
| Female | 6 | 10 |
| Age (weeks) | 45.5 (0.3) | 45.7 (0.3) |
| Body Mass (g) | 470 (32.9) | 464 (37.9) |
| Heart Mass (g) | 1.4 (0.1) | 1.3 (0.09) |
| Heart Mass to Body Mass (mg/g) | 3.1 (0.14) | 3.0 (0.16) |
| Heart Rate (beats/min) | 318.7 (8.8) | 327.2 (8.0) |
| Left Ventricular Width (mm) | 3.6 (0.14) | 3.5 (0.10) |
| Left Ventricular Width to Body Mass (mm/mg) | 8.0 (0.47) | 8.4 (0.61) |
| Septal Width (mm) | 3.2 (0.11) | 3.0 (0.08) |
| Septal Width to Body Mass (mm/mg) | 7.1 (0.49) | 7.0 (0.43) |
| Cardiac Output (mL/min) | 76.5 (3.7) | 82.4 (4.0) |
| Stroke Volume (μL) | 240 (10.0) | 253 (12.7) |
| Ejection Fraction (%) | 72 (3.6) | 77 (2.0) |
| Tibialis Anterior Mass (g) | 1.6 (0.11) | 1.5 (0.11) |
| Total hippocampus wet weight (g) | 0.11 (0.003) | 0.11 (0.002) |
| Right hippocampus wet weight (g) | 0.057 (0.001) | 0.056 (0.001) |
| Left hippocampus wet weight (g) | 0.058 (0.001) | 0.057 (0.001) |
| Right hippocampal Citrate Synthase Activity<br>(μmol/g/min) | 31.5 (5.0) | 31.1 (4.3) |
| Left hippocampal Citrate Synthase Activity<br>(μmol/g/min) | 36.4 (4.3) | 28.1 (3.0) |

**Note:** TAC = transverse aortic constriction, SHAM = sham control, g = grams, mg/g = milligrams per gram, beats/min = beats per minute, mm = millimeter, mm/mg = millimeter per milligrams, mL/min = milliliter per minute, μL = microliter, % = percent, μmol/g/min = micromole per gram per minute. Data presented as means (M) and standard error of the mean (SEM).

## Results

### Animal Characteristics

Thirty-one animals were utilized (7 TAC male, 6 TAC female, 7 SHAM male, and 10 SHAM female) in this study. Animal characteristics for the full (n=31) cohort can be found in Table 1. Overall, no significant differences between TAC and SHAM were observed in any of the characteristics, including hippocampal mitochondrial content, heart mass, septal width, cardiac output, ejection fraction, and stroke volume (**Table 1**).

### TAC Carotid Blood Flow Velocities and Pulsatility were Chronically Lowest on the Left Hemisphere and Highest on the Right Hemisphere

At all the time points measured, 2×2 ANOVA revealed there were significant main effects and interactions on peak carotid blood flow velocity (**Fig. 1B**, **Table S1**) and pulsatility index (**Fig. 1C, Table S2**). At 20 weeks, peak artery velocity was lower in the left carotid of TAC (M= 902.1 ± 99.4) when compared to the right (M=1716.8 ± 84.9 mm/sec, *p* <0.001) of TAC, as well as the right (M=1450.5 ± 59.1 mm/sec, *p* <0.001) and left (M= 1376.3 mm/sec ± 73.9, *p* <0.01) of SHAM animals (**Fig. 1B**). These patterns held steady at 30- and 40 weeks post-surgery, and peak blood flow velocity remained lowest in the left carotid of TAC (30 weeks M= 898.5 ± 129.2 mm/sec, 40 weeks M= 794.9 ± 103.6 mm/sec) when compared to the right of TAC (30 weeks M=1692.0 ± 85.2 mm/sec, *p* <0.001, 40 weeks M=1573.7 ± 81.9 mm/sec, *p* <0.001), as well as the right (30 weeks M= 1490.8 ± 81.2 mm/sec, *p* <0.01, 40 weeks M= 1551.7 ± 131.2 mm/sec, *p* <0.01) and left (30 weeks M= 1505.0 ± 92.9 mm/sec, *p* <0.001, 40 weeks M= 1442.0 ± 85.7 mm/sec, *p* <0.001) of SHAM animals (**Fig. 1B**). This hemodynamic phenotype did not worsen as animals aged as there were no significant main effects of Time (**Table S3**) on carotid artery velocities or pulsatility.

Blood flow pulsatility was consistently highest on the right carotid of TAC animals while simultaneously lowest on the left carotid. Indeed, at 20 weeks post-surgery, blood flow pulsatility was significantly higher in the right carotid of TAC (M=1.43 ± 0.02 P.I) animals when compared to the left (M=1.05 ± 0.04 P.I, *p* <0.001) of TAC, as well as the right (M= 1.30 ± 0.03 P.I, *p* <0.05) and left (M= 1.23 ± 0.02 P.I, *p* <0.001) of SHAM animals (**Fig. 1C**). Carotid blood flow pulsatility remained the highest on the right carotid of TAC at all other time points (30 weeks M= 1.39 ± 0.03 P.I, 40 weeks M= 1.43 ± 0.03 P.I) when compared to the left (30 weeks M = 0.98 ± 0.05 P.I, *p*<0.001, 40 weeks M= 0.86 ± 0.04 P.I, *p* <0.001) of TAC, as well as the right (30 weeks, M= 1.28 ± 0.02, p >0.05, 40 weeks M= 1.29 ± 0.03 P.I, *p* <0.05) and left (30 weeks M= 1.22 ± 0.02 P.I, *p*<0.01, 40 weeks M= 1.22 ± 0.02 P.I, *p* <0.001) carotids of SHAM controls (**Fig. 1C**).

Lastly, carotid flow rate was also calculated as previously described ^33^ and was found to be lower in the left (20 weeks M= 0.58 ± 0.07, 30 weeks M= 0.59 ± 0.1, 40 weeks M= 0.51 ± 0.07) of TAC animals at all time points when compared to the right (20 weeks M= 1.06 ± 0.07, *p* <0.001, 30 weeks M= 1.07 ± 0.05, *p* < 0.001, 40 weeks M=1.02 ± 0.05, *p*<0.001) of TAC, as well as the right (20 weeks M= 0.90 ± 0.04, *p* <0.01, 30 weeks M= 0.91 ± 0.04, *p* < 0.05, 40 weeks M= 0.98 ± 0.06, *p*<0.001) and left (20 weeks M = 0.84 ± 0.04, *p* <0.05, 30 weeks M = 0.91 ± 0.06, *p* < 0.05, 40 weeks M = 0.94 ± 0.05 *p*<0.001) of SHAM (**Fig. 1D**). No significant main effects of time were observed on either blood flow pulsatility (**Table S2**) or carotid flow rate.

Sex differences in TAC animals were only observed on the right side, and females tended to showcase significantly greater peak velocities at 30-, and 40 weeks than males (**Fig. S2A**). No significant sex differences were seen in right carotid blood flow pulsatility in TAC (**Fig. S2A**). In SHAM animals, females displayed significantly greater left and right peak blood flow velocities at 40-weeks than males (**Fig. S2B**). When compared to male SHAM, blood flow pulsatility was also found to be significantly higher in female SHAM animals in the left at 20-, and 30 weeks post-surgery (**Fig. S2B**) and at 20- and 40 weeks on the right-side (**Fig. S2B**).

### Carotid Artery Cross-Sectional Area and Diameter are Influenced by Sex but not TAC Surgery

Carotid artery cross sectional area in systole was calculated as: CSA = D^2^ × 0.785 ^34^, and no significant main effects or interactions on systolic cross-sectional area were detected (**Fig. S3A**). Independently run 2×2 ANOVA models revealed no significant main effects nor Condition by Hemisphere interactions on either systolic or diastolic carotid artery diameter at any of time points measured (**Fig. S4A, & S5A**). There were also no significant main effects of Time on either systolic carotid artery diameter, diastolic artery diameter (**Table S4**), nor systolic cross-sectional area in either TAC or SHAM animals.

As it pertains to sex differences, TAC females had on average significantly lower cross-sectional area, as well as systolic and diastolic artery diameters than males, but these differences were only significant on the right side at 20 weeks (**Fig. S3B, S4B, & S5B**). In SHAM animals, females displayed significantly lower systolic cross-sectional area on the right and left side at most time points when compared to males (**Fig. S3C**). Further, sex differences in carotid artery diameter were extensive, and on average, females displayed lower left systolic and diastolic diameters than males at all time points (**Fig. S4C & S5C**). On the right side, females also had significantly lower systolic and diastolic diameters than males at most time points (**Fig. S4C & S5C**).

### Hippocampal Mitochondrial Respiration is Impaired in the Right, but not the Left, Hemisphere of TAC Animals

A significant main effect of Hemisphere (*F*(1,56)=5.08, *p*<0.05, ηp^2^=0.08) in CI-linked coupled respiration was observed and follow up contrast revealed that within condition hemispheric differences were greatest in TAC (*t*(56)=1.94, *p*=0.057) than SHAM (*t*(56)=1.18, *p*=0.24), but these hemispheric differences within TAC did not reach statistical significance (*p=*0.057) (**Fig. 2A**). There was also a significant Condition by Hemisphere interaction (*F*(1,56)=4.69, *p*<0.05, ηp^2^=0.077) for CII-linked uncoupled respiration and follow up contrasts within TAC revealed that the right hippocampus had significantly lower CII-linked uncoupled respiration when compared to the left hippocampus (*t*(56)=-2.60, *p*=0.012) (**Fig. 2A**). Lastly, because there was a main effect of Hemisphere that approached significance (*F*(1,56)=3.75, *p*=0.058, ηp^2^=0.063) for CI&II-linked coupled respiration, this was followed up with a contrast analysis, and the right hippocampus of TAC animals had significantly lower respiration rates than the left (*t*(56)=-2.29, *p*=0.025) (**Fig. 2A**). No main effects of Condition (*F*(1,56)=1.06, *p*=0.30, ηp^2^=0.01), Hemisphere (*F*(1,56)=0.055, *p*=0.81, ηp^2^=0.0009), nor Condition by Hemisphere interaction (*F*(1,56)=0.88, *p*=0.35, ηp^2^=0.01) were observed for mitochondrial content as measured via citrate synthase activity (**Table 1**).

**Figure 2.**
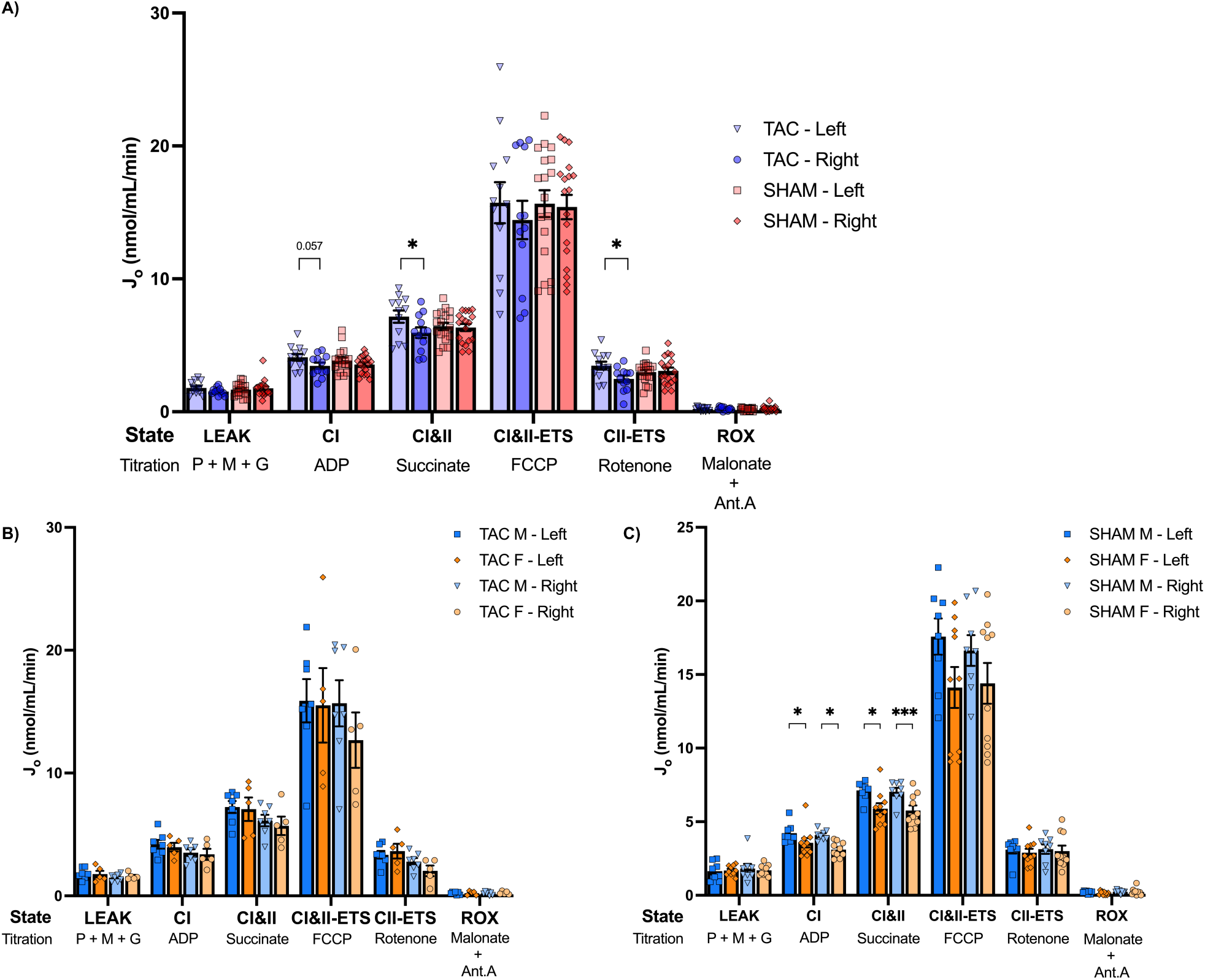
Hippocampal mitochondrial respiration is impaired in the right hemisphere in TAC, and SHAM animals display sex dimorphisms not present in TAC: **A)** Hippocampal mitochondrial respiration in right and left hemisphere of TAC and SHAM Animals. **B)** Right and left hemispheric hippocampal. Mitochondrial respiration in male and female TAC. **C)** Right and left hemispheric hippocampal. Mitochondrial respiration in male and female SHAM. J_O_ = oxygen consumption, nmol/mL/min = nanomoles per milliliter per minute, LEAK-leak respiration, CI = complex I, CII = complex II, ETS = electron transfer system, ROX = residual oxygen consumption rate, P = pyruvate, M = malate, G = glutamate, TAC = transverse aortic constriction, SHAM = sham controls, M = male, F = female. Data presented as mean + SEM, * = p<0.05. Data presented as means + SEM.

No sex differences in mitochondrial respiration were found in TAC animals (**Fig. 2B**). However, in SHAM controls, CI-linked coupled respiration in female SHAM was lower than when compared to male counterparts in both the right (M=3.10±0.16 v. M=4.09±0.12, *p*<0.05, d=-2.22) and left (M=3.57±0.31v. M=4.22±0.23, *p*<0.05, r=-0.68) hippocampus (**Fig. 2C**). Female SHAM also displayed lower CI&II-linked coupled respiration when compared to male counterparts in both the right (M=5.76±0.33v. M=7.04±0.26, *p*<0.05, r=-0.71) and the left (M=5.88±0.37 v. M=7.12±0.22, *p*<0.05, d=-1.27) hippocampus (**Fig. 2C**).

Pearson product-moment correlations were used as an exploratory analysis to determine if carotid artery blood flow velocities and pulsatility 40 weeks post-surgery were related to mitochondrial respiration. For this analysis, only SHAM and TAC animals with a complete set of hemodynamic data were included. That is, the 3 TAC animals that were found to have poor left carotid time series were excluded from this analysis. Overall, carotid artery peak velocity (*r* = −0.453, *p*= 0.004, [−0.67, −0.16]) (**Fig. 3A**), blood flow pulsatility (*r* = −0.429, *p*= 0.006, [−0.65, −0.13]) (**Fig. 3B**), and mean velocity (*r* = −0.448, *p*= 0.004, [−0.66, −0.15]) (**Fig. S6A**) were all negatively associated to CI&II-linked coupled respiration, and higher velocities and pulsatility were associated with impaired respiration rates. Similarly, carotid artery peak velocity (*r* = −0.473, *p*= 0.002, [−0.68, −0.18]) (**Fig. 3C**), and blood flow pulsatility (*r* = −0.481, *p*= 0.002, [−0.69, −0.19]) (**Fig. 3D**), and mean velocity (*r* = −0.45, *p*= 0.003, [−0.67, −0.16]) (**Fig. S6B**) were also negatively associated to CII-linked uncoupled respiration, and higher velocities and pulsatility were associated with lower uncoupled respiration rates.

**Figure 3.**
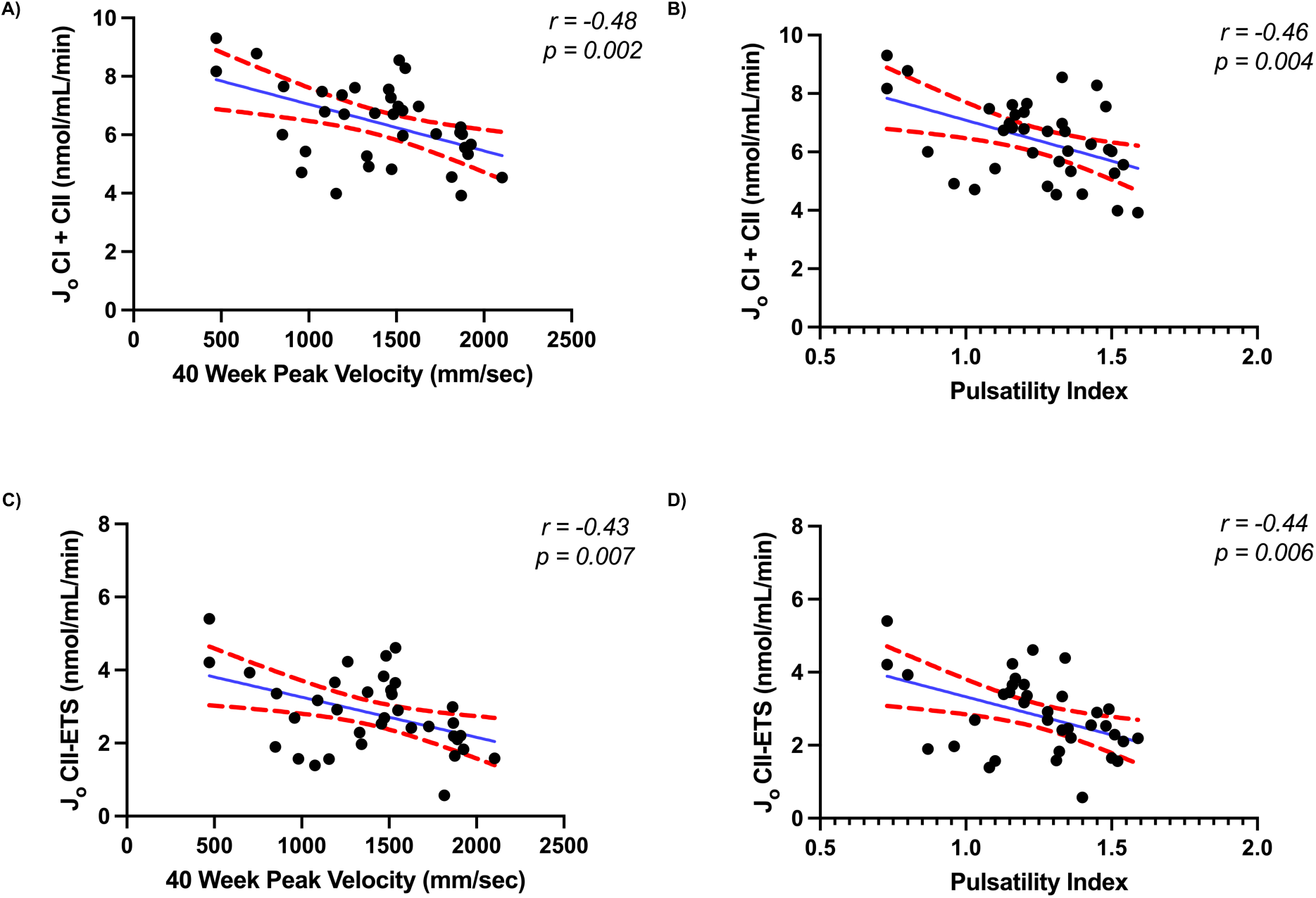
High carotid blood flow velocity and pulsatility are negatively correlated with hippocampal mitochondrial respiration: **A)** Peak carotid artery velocity and **B)** pulsatility index are negatively associated with CI+II coupled hippocampal respiration. **C)** Peak Carotid artery velocity and **D)** pulsatility index are negatively associated with CII uncoupled hippocampal respiration. J_O_ = oxygen consumption rate, nmol/ml/min = nanomoles per milliliter per minute, CI = complex I, CII = complex II, ETS = electron transfer system.

### Brain Hypoperfusion leads to an Upregulation in Protein Expression of Mitochondrial Fusion Proteins

There were significant main effects of Condition for both the long- (*F*(1,56)=4.92, *p*<0.05, ηp^2^=0.081) and short- (*F*(1,56)=5.01, *p*<0.05, ηp^2^=0.082) (**Fig. 4A**) OPA1 isoforms, as well as total OPA1 (*F*(1,56)=5.09, *p*<0.05, ηp^2^=0.083) (calculated as the sum of the density signal of the short- and long-isoforms) with TAC animals displaying higher protein expression than SHAM animals. Further, a significant main effect of Condition (*F*(1,56)=6.53, *p*<0.05, ηp^2^=0.10), where TAC animals showcased higher protein expression than SHAM was also detected for HSP-60 (**Fig. 4A**). No significant main effects of Condition, Hemisphere, nor Condition by Hemisphere interaction were detected for MFN2 (**Fig. 4A**), FIS1, nor DRP1 protein expression (**Fig. 5A**).

**Figure 4.**
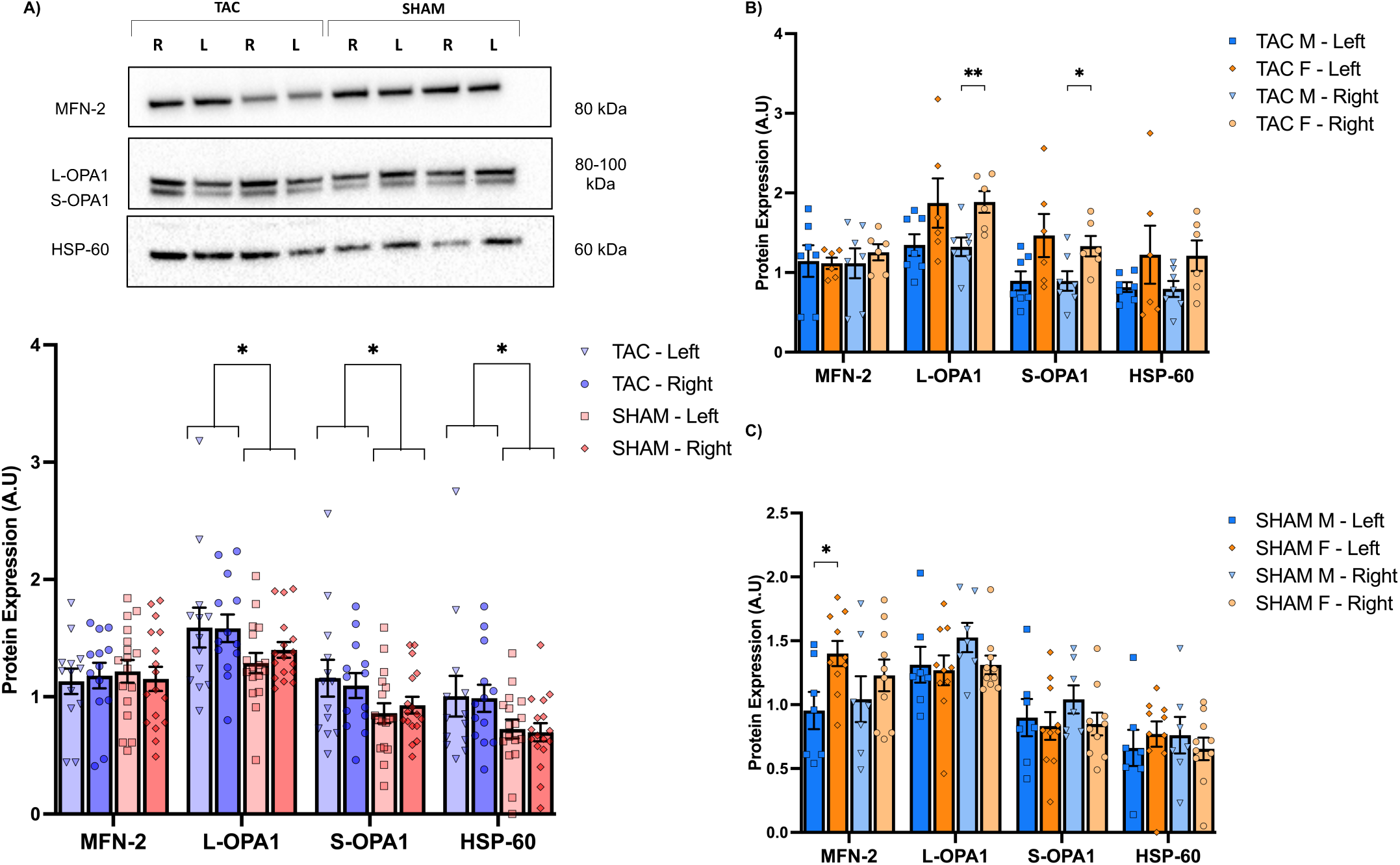
Protein expression of hippocampal mitochondrial fusion markers is higher in TAC compared to SHAM rats. **A)** Protein expression of mitochondrial MFN-2, L-OPA1, S-OPA1, and HSP-60 in the right and left hippocampus of TAC compared to the right and left hippocampus of SHAM rats. **B)** Protein expression of mitochondrial fusion markers in male compared to female TAC. **C)** Protein expression of mitochondrial fusion markers in male compared to female SHAM. All protein expression is normalized to total protein. A.U = arbitrary units, MFN = mitofusin 2, L-OPA1 = Optic Atrophy 1 long isoform, S-OPA1 = Optic Atrophy 1 short isoform, HSP-60 = heat shock protein-60, TAC = transverse aortic constriction, SHAM = sham controls, M = male, F = female, kDa = kilodalton, * = p<0.05, ** = p<0.01. Data presented as means + SEM.

**Figure 5.**
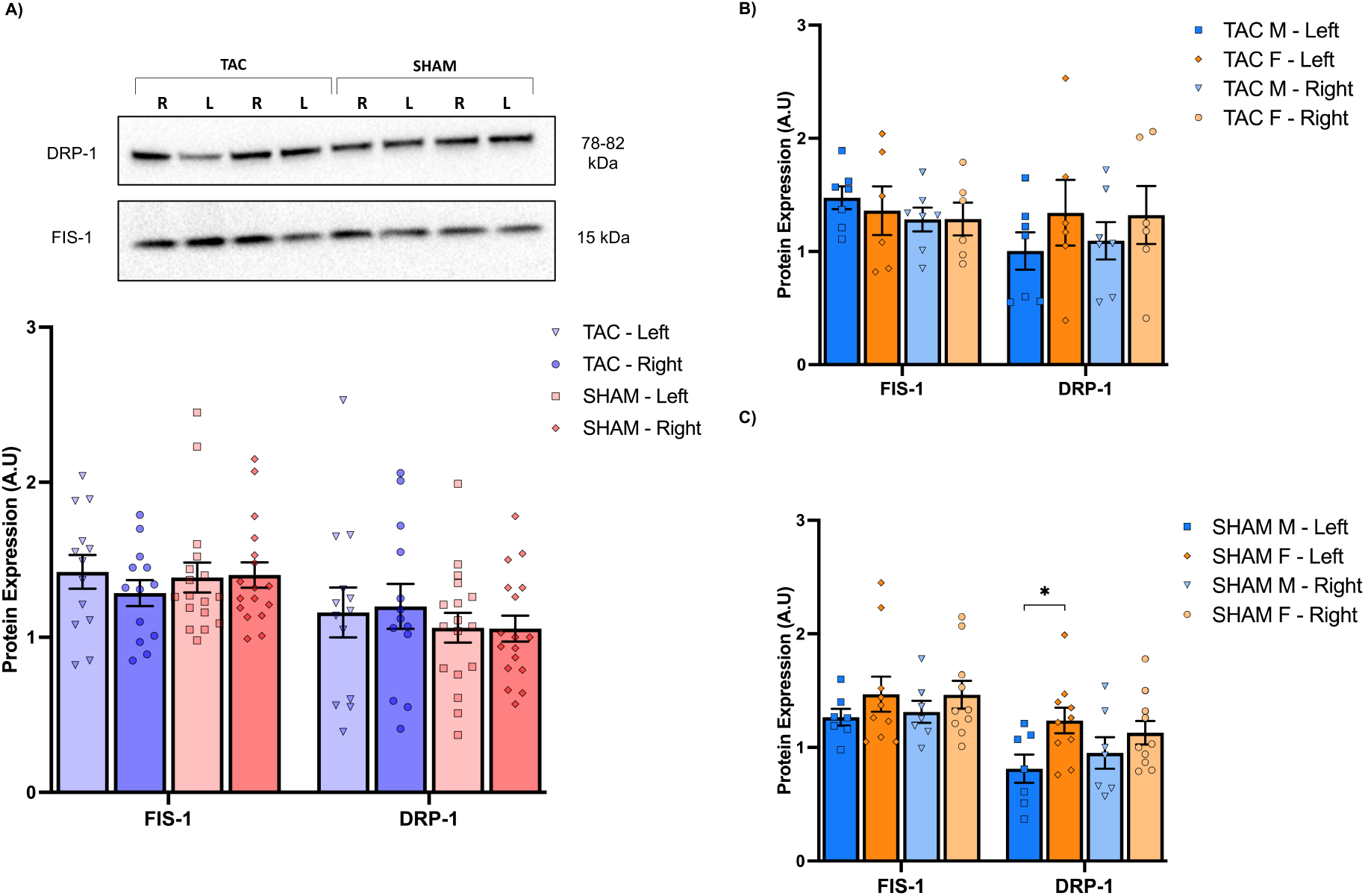
Hippocampal protein expression of mitochondrial Fission Markers are not Altered in TAC compared to SHAM, and Fis1 is higher in sham females compared to sham males. **A)** Protein expression of mitochondrial FIS-1 and DRP-1 in TAC and SHAM rats. **B)** FIS-1 and DRP-1 protein expression in male compared to female TAC animals. **C)** FIS-1 and DRP-1 protein expression in male compared to female SHAM animals. Individual protein expression was normalized to total protein. A.U = arbitrary units, FIS1 = fission 1, DRP1 = dynamin-related protein 1, TAC = transverse aortic constriction, SHAM = sham controls, M = male, F = female, kDa = kilodalton, * = p <0.05. Data presented as means + SEM.

In TAC animals, sex differences in protein expression were only found within the right hippocampus, where females were found to have significantly higher protein expression of the short (M=1.33±0.12 v. M=0.89±0.12, *p*<0.05, d=1.36) and long (M=1.88±0.13 v. M=1.32±0.11, *p*<0.05, d=1.76) OPA1 isoforms compared to males (**Fig. 4B**). No significant sex differences in hippocampal protein expression were found in the left hemisphere of TAC. In SHAM animals, sex differences in expression of quality control proteins were found on the left hippocampus and on average, females had higher protein expression than males of MFN2 (M=1.40±0.22 v. M=0.95±0.40, *p*<0.05, d=1.30) (**Fig. 4C**), and DRP1 (M=1.23±0.11 v. M=0.81±0.12, *p*<0.05, d=1.22) (**Fig. 5C**). No other sex differences were evident in SHAM controls.

### Protein Expression of Glucose Transporters in Unaltered in TAC Animals

Overall, we found no significant main effects of Condition (*F*(1,56)=0.424, *p*=0.51, ηp^2^=0.008), Hemisphere (*F*(1,56)=0.33, *p*=0.56, ηp^2^=0.006), nor Condition by Hemisphere interaction (*F*(1,56)=0.11, *p*=0.73, ηp^2^=0.002) in hippocampal protein expression of GLUT-1 (**Fig. 6**). Similarly, no significant no main effects of Condition (*F*(1,56)=0.38, *p*=0.53, ηp^2^=0.007), Hemisphere (*F*(1,56)=0.22, *p*=0.63, ηp^2^=0.004), nor a Condition by Hemisphere interaction (*F*(1,56)=0.09, *p*=0.75, ηp^2^=0.002) for hippocampal protein expression of GLUT-3 (**Fig. 6**). Further, no sex differences were found in the protein expression of glucose transporters in either TAC or SHAM.

**Figure 6.**
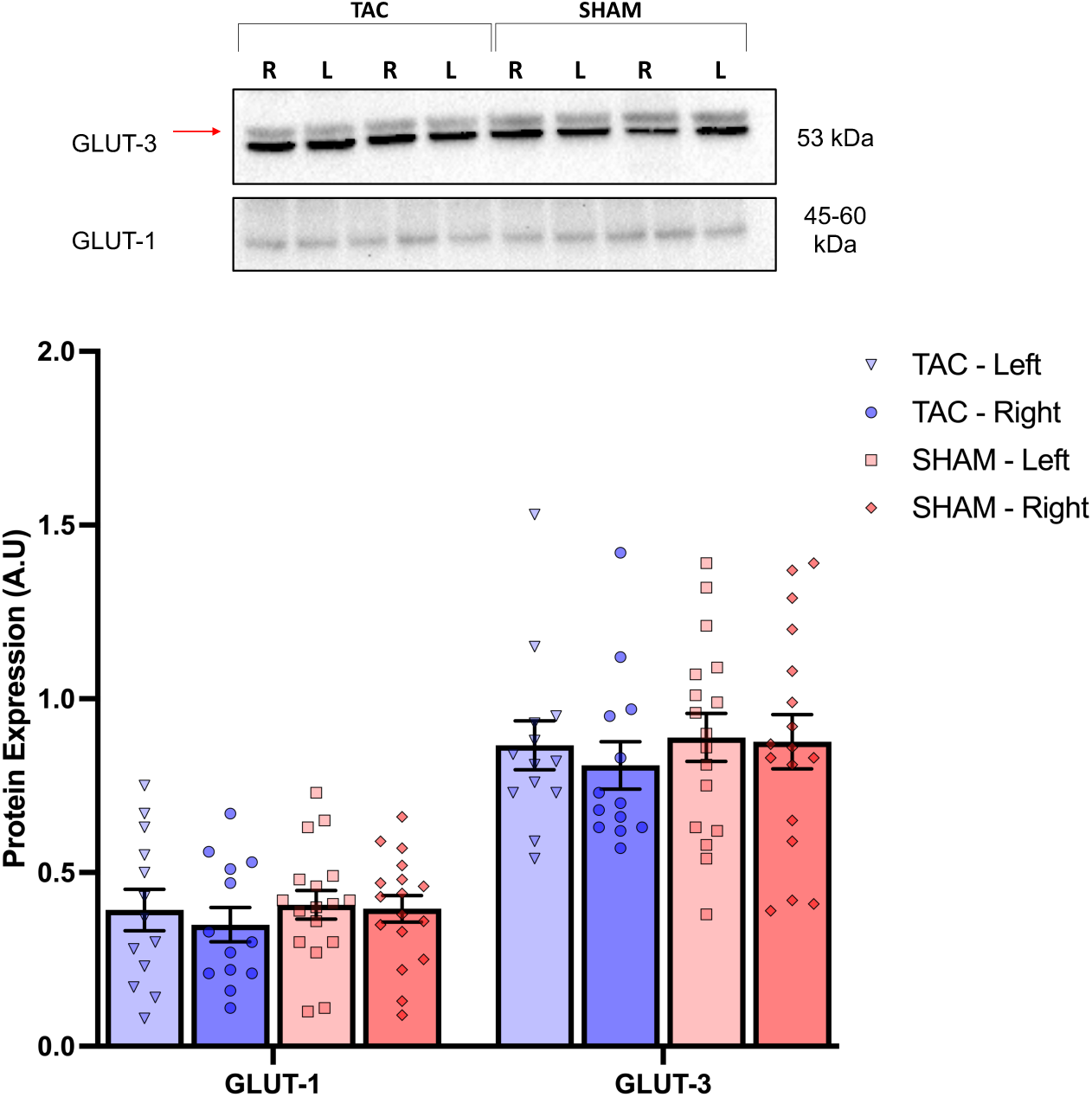
Protein Expression of Glucose Transporters is not Altered in TAC: Glut-3 and Glut-1 protein expression in left and right hippocampus of TAC and SHAM rats, normalized to total protein. A.U = arbitrary units, GLUT-1 = Glucose Transporter 1, GLUT-3 = Glucose transporter 3, TAC = transverse aortic constriction, SHAM = sham controls, kDa = kilodalton. Data presented as means + SEM.

## Discussion

While primarily employed to cause cardiac pressure overload, transverse aortic constriction also alters blood flow within the carotid arteries so that right carotid artery (upstream of the constriction) blood flow velocities and pulsatility are greater than the left (downstream of the constriction). Thus, the transverse aortic constriction model is uniquely suited to simultaneously study how the two primary hemodynamic mechanisms behind brain hypoperfusion^16, 35^ influence cellular processes associated with neurodegeneration. Overall, we report distinct carotid hemodynamics are coupled to hemispheric differences in coupled and uncoupled mitochondrial respiration, where the right hippocampus shows deficits in mitochondrial respiration and the left hippocampus does not. We also report TAC animals have higher expression of mitochondrial fusion and uncoupled protein response (UPR^MT^) proteins in both hemispheres, compared to SHAM animals. Together, these findings suggest the molecular mechanisms linking cardiovascular disease to neurodegeneration could be uniquely influenced by perfusion patterns and could inform future studies on flow-dependent processes.

### Carotid Artery Hemodynamics

Carotid blood flow velocities are higher in the right carotid of TAC animals when compared to the left carotid at every time point following constriction, and carotid blood flow pulsatility is higher in the right of TAC animals when compared to not only the left carotid within TAC, but also the right and left carotids of SHAM animals. This is in line with existing literature showing arterial pulse pressure is highest in the right carotid of TAC animals when compared to the right and left of SHAM animals^26^ and may indicate losses in arterial compliance are greatest in the right carotid of TAC. It has been previously reported that the divergent blood flow phenotype in TAC can develop as soon as 1 day post-surgery and remain largely unchanged 7 days post-surgery^27^. A similar pattern has been reported in a mouse model of Alzheimer’s Disease with TAC-induced brain hypoperfusion, where the bilateral hemodynamic phenotype is present 3 weeks post-TAC, and largely unchanged 8 weeks post-TAC^36^. Because most studies assessing carotid artery hemodynamics in TAC have employed acute (1-day, and 7-day post-TAC)^27^, or pre-and post-models^26, 36, 37^, there is little data on characterizing carotid blood flow in TAC from a chronic point of view. Further, these rats were 20-weeks old at the first ultrasound assessment, and therefore growth was unlikely to contribute to worsening hemodynamics. As such, these results build on the existing literature and show TAC carotid flow characteristics are stable over time and can be therefore adapted for chronic studies.

Recent work by De Montgolfier and colleagues provides compelling evidence showing that, 6 weeks post-TAC surgery, leads to two distinct hemodynamic phenotypes that induce hypoperfusion, yet the pulsatility associated with the right carotid leads to a more pronounced degree of hypoperfusion in the right cerebral hemisphere when compared to the left^26^. Moreover, these TAC-induced hemispheric differences in brain perfusion are accompanied by other hemispheric differences such as greater deficits in blood brain barrier integrity, higher adverse microvascular events (e.g., microbleeds), and greater losses in microvascular density in the right hemisphere when compared to the left^26^. Thus, although blood flow velocity in the right carotid of TAC does not substantially differ to SHAM controls in this study, the higher sustained pulsatility seen in the right carotid of TAC may be accompanied by sustained hypoperfusion, cerebral microvascular stress, and neurovascular damage. Overall, this longitudinal characterization of carotid blood flow velocity and pulsatility resulting from TAC demonstrates this model may be suitable to study not only brain hypoperfusion, but also how the mechanisms that give rise to hypoperfusion^26^ influence various neurophysiological processes such as substrate transport and metabolism.

Surprisingly, the impact of TAC on left and right carotid artery velocities and pulsatility were not affected by time. While surprising at first, in acute time windows, the TAC hemodynamic phenotype has been reported to have a rapid onset with little change after being developed^27, 36^

Although our TAC animals tend to display a phenotype suggestive of impaired cardiac function and cardiac hypertrophy, these are not significantly different than our SHAM controls. Indeed, the model was intended to produce a modest cardiac phenotype and includes both male and female animals, increasing variability in size-dependent parameters. To our knowledge, we are the first to extend the TAC model in rats to 40 weeks post-surgery, and the use of a 20G needle facilitated the survivability of our animals to the intended end point. Moreover, we note that while we do not have a robust heart failure phenotype, our TAC animals display the same carotid hemodynamic phenotype reported in previous work with aggressive ligation aproaches^23, 26, 27, 36^. Therefore, the blunted cardiac function and cardiac hypertrophy phenotypes did not translate to a loss in the brain hemodynamics that are central to this study.

### Mitochondrial Respiration and Quality Control

Mitochondria are key organelles that supply the vast majority of ATP utilized by neurons^15, 38–40^ and may be key organelles compromised in chronic brain hypoperfusion. A primary finding of this study is hemispheric differences in mitochondrial respiration following TAC. Specifically, when compared to the left hippocampus, the right hippocampus of TAC animals displays significantly lower CI&II-linked coupled respiration, as well as CII-linked uncoupled respiration. Further, despite not reaching statistical significance (*p*=0.057), we note that the right hippocampus in TAC also displays lower CI-linked coupled respiration when compared to the left hippocampus (M=3.46±0.23 v M=4.10±0.24) which may be of physiological relevance. Because coupled respiration represents the maximal respiratory capacity of the electron transfer pathways in the presence of saturating substrates and ADP^41^, CI&II-linked coupled respiration represents the upper limits in coupling the transfer of electrons along the electron transport system (ETS) to the phosphorylation of ADP to ATP^42^. Moreover, because uncoupled respiration employs protonophores to disrupt the inner mitochondrial membrane and dissipate the protonmotive force, CII-uncoupled represents the maximal electron transfer through complex II, III, & IV^41, 43^. Thus, the deficits in coupled and uncoupled respiration in the right hippocampus may suggest that high pulsatility may have effects that propagate into cellular organelles like the mitochondria and disrupt the structural and functional properties of the ETS. Moreover, because previous studies show TAC induces a more robust hypoperfusive response in the right hemisphere than the left^26^, our results compliment these observations and show the metabolic consequences of hypoperfusion are also more pronounced on the right hemisphere than the left.

Meanwhile, despite no hemispheric differences in the protein expression mitochondrial quality control markers in TAC, markers associated with mitochondrial fusion and the UPR^MT^ were upregulated in both hemispheres of TAC animals when compared to SHAM controls. Both the long- and short-OPA1 isoforms, as well as HSP-60 are significantly higher in hippocampus of TAC compared to SHAM. It is worth noting that: 1) the UPR^MT^ and HSPs are integral for the regulation of ETC stoichiometries during mitochondrial remodeling and complex formation^39, 44^, 2) inner-membrane bound OPA1 is integral for the regulation of mitochondrial cristae shape^45^, and 3) whole brain mitochondrial complex assembly formation is impaired in chronic hypertension^24^.

Therefore, it is possible that the bilateral upregulation of OPA1 in TAC may be an adaptive response to increase mitochondrial cristae complexity to improve metabolism in the presence of low tissue perfusion, while the dual upregulation of HSP-60 may serve to ensure mitochondrial stoichiometries and ETC complex formation are sustained and proteotoxic stress is minimized. However, this possibility should be better characterized in future studies, and high-resolution imaging of mitochondria cristae, as well as the roles of other prominent HSPs such as mortalin and TRAP-1 should all be addressed. Nevertheless, taken collectively, these results suggest neural mitochondria may be able to upregulate quality control mechanisms in the presence of brain hypoperfusion, but these adaptive responses may be insufficient when hypoperfusion stems from high blood flow pulsatility.

### Glucose Transporter Protein Expression

Surprisingly, there are no hemispheric differences in the expression of either GLUT-1 or GLUT-3 within TAC, nor any differences between TAC and SHAM. Poulet and colleagues report hippocampal GLUT-1 (but not GLUT-3) mRNA is downregulated in TAC animals when compared to SHAM controls 3 weeks and 4 weeks post-surgery^36^. Intriguingly, when looking at the cortex, GLUT-1 was only downregulated 4 weeks post-surgery^36^. Because ischemia reperfusion studies have shown acute increases in blood flow are accompanied by an initial upregulation in GLUT mRNA levels^46^, it is possible that downregulation of GLUT mRNA in acute models of TAC is also a temporary response^47, 48^.

### Sex Differences

Historically, cardiovascular disease has been studied in the context of male physiology and disease^49^, despite the fact that the prevalence of cardiovascular disease is greater in older women than older men^50^. We therefore performed an exploratory analysis to determine whether there are sex differences in right and left carotid hemodynamics, mitochondrial respiration, and protein expression of target markers. Blood flow velocity is greater in females compared to male SHAM animals at the 40-week timepoint, and these sex differences occur earlier in TAC (30-, and 40-weeks), with higher blood flow velocities in the right side of females. Because the right side of TAC showcased the sex differences in blood flow velocities 10 weeks earlier than SHAM controls, this may suggest that higher blood flow velocities and associated pulsatility may speed up the development of sex dimorphisms in vascular health. In support of this interpretation, recent frameworks show the cerebral vasculature of non-diseased female Sprague-Dawley rats has different structural properties than male counterparts such as lower content of vascular smooth muscle cells, higher collagen deposits, and thicker internal elastic lamina^51^. These structural differences, in turn, seem to impair the contractile capabilities of large cerebral arteries, lower arterial distensibility, elevate the myogenic response, and have negative effects on the vascular functional capabilities in females when compared to males^51^. As such, female animals may have a reduced ability to cope with sustained higher pulsatile blood flow velocities over time, and the early manifestation of sex dimorphisms in blood flow seen in TAC could signify an early failing vasculature.

As it pertains to hippocampal mitochondrial physiology, SHAM females have significantly lower CI-linked and CI&II-linked coupled respiration, concomitant with higher protein expression of fusion (MFN2) and fission (DRP1) markers compared to SHAM males. These observations contribute to existing animal data where sex-based differences in mitochondrial function and quality control are reported. For example, CI-linked respiration is higher in cortical mitochondrial fractions obtained from 13-month old female mice compared to males^52^. Hormones such as estradiol may also influence brain mitochondria^53^, as complete removal of estrogens in ovariectomized macaque monkeys is reported to alter brain mitochondrial shape from elongated tubules to a more donut-shaped phenotype^54^, which in this context, is associated with mitochondrial stress^53^. Thus, the presence of sex dimorphisms in respiration and markers of mitochondrial dynamics in our SHAM animals could suggest that circulating hormone levels may be partially involved, but this should be addressed further in future studies. Nevertheless, our results contribute to a growing field of research and may aid in better understanding how sex dimorphisms in brain mitochondrial function and dynamics change throughout the lifespan and how these relate to cardiovascular health and disease.

### Limitations

While this study did assess carotid blood flow velocity and pulsatility, it did not quantify hemispheric or hippocampal tissue perfusion. As such, the data from this study cannot directly show that altered carotid blood flow in TAC led to brain hypoperfusion. However, previous work has established that TAC does in fact lead to whole brain hypoperfusion and that this hypoperfusion is not the same between the two cerebral hemispheres ^23, 27^. Thus, because our animals displayed the same phenotype observed in prior TAC hypoperfusion studies, it is also likely that our TAC animals did develop brain hypoperfusion. Additionally, this study did not include the use of behavioral testing assays (such as the Morris Water Maze or Y-Maze) that are normally used to link adverse physiological changes to cognitive health. Molecular data and cognitive batteries are key in the establishment of animal models to study neurodegeneration and although there is limited evidence showing behavioral deficits in TAC ^26^ future work should aim to include cognitive assays to fully characterize the behavioral consequences of TAC and how they relate to the molecular changes that happen within cortical and subcortical structures.

## Conclusion

While transverse aortic constriction is traditionally used to model cardiac pressure overload, the present study highlights its utility for investigating flow-dependent mechanisms of brain hypoperfusion and their impact on neurophysiological processes associated with neurodegeneration. Specifically, TAC leads to hemispheric differences in mitochondrial respiration and overall changes in quality control markers in the hippocampus. These findings underscore how distinct perfusion patterns—such as high blood flow velocities and pulsatility—affect mitochondrial function differently, offering new insights into the cellular mechanisms linking cardiovascular pathology and neurodegeneration.

## Supplemental Material

Tables S1-S4

Figure S1-S6

## References

1. Le Couteur DG, Thillainadesan J. What is an aging-related disease? An epidemiological perspective. The Journals of Gerontology: Series A. 2022.

2. Niccoli T, Partridge L. Ageing as a risk factor for disease. Current biology. 2012; 22:R741–R52.

3. Ahmad FB, Anderson RN. The leading causes of death in the US for 2020. Jama. 2021; 325:1829–30.

4. Prince MJ, Wu F, Guo Y, Robledo LMG, O’Donnell M, Sullivan R, et al. The burden of disease in older people and implications for health policy and practice. The Lancet. 2015; 385:549–62.

5. de La Torre JC. Cardiovascular risk factors promote brain hypoperfusion leading to cognitive decline and dementia. Cardiovascular psychiatry and neurology. 2012; 2012.

6. Association As. 2019 Alzheimer’s disease facts and figures. Alzheimer’s & Dementia. 2019; 15:321–87.

7. T O’Brien J, Thomas A. Vascular dementia. The Lancet. 2015; 386:1698–706.

8. Rajeev V, Fann DY, Dinh QN, Kim HA, De Silva TM, Lai MK, et al. Pathophysiology of blood brain barrier dysfunction during chronic cerebral hypoperfusion in vascular cognitive impairment. Theranostics. 2022; 12:1639.

9. Du S-Q, Wang X-R, Xiao L-Y, Tu J-F, Zhu W, He T, et al. Molecular mechanisms of vascular dementia: what can be learned from animal models of chronic cerebral hypoperfusion? Molecular neurobiology. 2017; 54:3670–82.

10. Kalaria RN. The pathology and pathophysiology of vascular dementia. Neuropharmacology. 2018; 134:226–39.

11. Shabir O, Berwick J, Francis SE. Neurovascular dysfunction in vascular dementia, Alzheimer’s and atherosclerosis. BMC neuroscience. 2018; 19:1–16.

12. Sabayan B, Jansen S, Oleksik AM, van Osch MJ, van Buchem MA, van Vliet P, et al. Cerebrovascular hemodynamics in Alzheimer’s disease and vascular dementia: a meta-analysis of transcranial Doppler studies. Ageing research reviews. 2012; 11:271–7.

13. Haller S, Zaharchuk G, Thomas DL, Lovblad K-O, Barkhof F, Golay X. Arterial spin labeling perfusion of the brain: emerging clinical applications. Radiology. 2016; 281:337–56.

14. Watts JM, Whitlow CT, Maldjian JA. Clinical applications of arterial spin labeling. NMR in Biomedicine. 2013; 26:892–900.

15. Cunnane SC, Trushina E, Morland C, Prigione A, Casadesus G, Andrews ZB, et al. Brain energy rescue: an emerging therapeutic concept for neurodegenerative disorders of ageing. Nature Reviews Drug Discovery. 2020; 19:609–33.

16. Ciacciarelli A, Sette G, Giubilei F, Orzi F. Chronic cerebral hypoperfusion: An undefined, relevant entity. Journal of clinical neuroscience. 2020; 73:8–12.

17. Meng L, Hou W, Chui J, Han R, Gelb AW. Cardiac output and cerebral blood flow: the integrated regulation of brain perfusion in adult humans. Anesthesiology. 2015; 123:1198–208.

18. Thorin-Trescases N, de Montgolfier O, Pinçon A, Raignault A, Caland L, Labbé P, et al. Impact of pulse pressure on cerebrovascular events leading to age-related cognitive decline. American Journal of Physiology-Heart and Circulatory Physiology. 2018; 314:H1214–H24.

19. Heiss W-D. PET imaging in ischemic cerebrovascular disease: current status and future directions. Neuroscience bulletin. 2014; 30:713–32.

20. Borja AJ, Hancin EC, Zhang V, Koa B, Bhattaru A, Rojulpote C, et al. Global brain glucose uptake on 18F-FDG-PET/CT is influenced by chronic cardiovascular risk. Nuclear Medicine Communications. 2021; 42:444–50.

21. Bordone MP, Salman MM, Titus HE, Amini E, Andersen JV, Chakraborti B, et al. The energetic brain–A review from students to students. Journal of neurochemistry. 2019; 151:139–65.

22. Venkat P, Chopp M, Chen J. Models and mechanisms of vascular dementia. Experimental neurology. 2015; 272:97–108.

23. Bink DI, Ritz K, Aronica E, Van Der Weerd L, Daemen MJ. Mouse models to study the effect of cardiovascular risk factors on brain structure and cognition. Journal of Cerebral Blood Flow & Metabolism. 2013; 33:1666–84.

24. Lopez-Campistrous A, Hao L, Xiang W, Ton D, Semchuk P, Sander J, et al. Mitochondrial dysfunction in the hypertensive rat brain: respiratory complexes exhibit assembly defects in hypertension. Hypertension. 2008; 51:412–9.

25. Du J, Ma M, Zhao Q, Fang L, Chang J, Wang Y, et al. Mitochondrial bioenergetic deficits in the hippocampi of rats with chronic ischemia-induced vascular dementia. Neuroscience. 2013; 231:345–52.

26. de Montgolfier O, Pinçon A, Pouliot P, Gillis M-A, Bishop J, Sled JG, et al. High systolic blood pressure induces cerebral microvascular endothelial dysfunction, neurovascular unit damage, and cognitive decline in mice. Hypertension. 2019; 73:217–28.

27. Poulet R, Gentile MT, Vecchione C, Distaso M, Aretini A, Fratta L, et al. Acute hypertension induces oxidative stress in brain tissues. Journal of Cerebral Blood Flow & Metabolism. 2006; 26:253–62.

28. Cauley E, Wang X, Dyavanapalli J, Sun K, Garrott K, Kuzmiak-Glancy S, et al. Neurotransmission to parasympathetic cardiac vagal neurons in the brain stem is altered with left ventricular hypertrophy-induced heart failure. American Journal of Physiology-Heart and Circulatory Physiology. 2015; 309:H1281–H7.

29. Henneman W, Sluimer J, Barnes J, Van Der Flier W, Sluimer I, Fox N, et al. Hippocampal atrophy rates in Alzheimer disease: added value over whole brain volume measures. Neurology. 2009; 72:999–1007.

30. Pena GS, Paez HG, Johnson TK, Halle JL, Carzoli JP, Visavadiya NP, et al. Hippocampal Growth Factor and Myokine Cathepsin B Expression following Aerobic and Resistance Training in 3xTg-AD Mice. International Journal of Chronic Diseases. 2020; 2020.

31. Burtscher J, Zangrandi L, Schwarzer C, Gnaiger E. Differences in mitochondrial function in homogenated samples from healthy and epileptic specific brain tissues revealed by high-resolution respirometry. Mitochondrion. 2015; 25:104–12.

32. Kuzmiak S, Glancy B, Sweazea KL, Willis WT. Mitochondrial function in sparrow pectoralis muscle. Journal of Experimental Biology. 2012; 215:2039–50.

33. Kenwright D, Thomson AJ, Hadoke P, Anderson T, Moran C, Gray G, et al. A protocol for improved measurement of arterial flow rate in preclinical ultrasound. Ultrasound International Open. 2015:E46–E52.

34. Blanco P. Volumetric blood flow measurement using Doppler ultrasound: concerns about the technique. Journal of ultrasound. 2015; 18:201–4.

35. Drouin A, Bolduc V, Thorin-Trescases N, Bélanger É, Fernandes P, Baraghis E, et al. Catechin treatment improves cerebrovascular flow-mediated dilation and learning abilities in atherosclerotic mice. American Journal of Physiology-Heart and Circulatory Physiology. 2011; 300:H1032–H43.

36. Carnevale D, Mascio G, Ajmone-Cat MA, D’Andrea I, Cifelli G, Madonna M, et al. Role of neuroinflammation in hypertension-induced brain amyloid pathology. Neurobiology of Aging. 2012; 33:205.e19-.e29.

37. Li H, Guo Q, Inoue T, Polito VA, Tabuchi K, Hammer RE, et al. Vascular and parenchymal amyloid pathology in an Alzheimer disease knock-in mouse model: interplay with cerebral blood flow. Molecular neurodegeneration. 2014; 9:1–15.

38. Raefsky SM, Mattson MP. Adaptive responses of neuronal mitochondria to bioenergetic challenges: Roles in neuroplasticity and disease resistance. Free Radical Biology and Medicine. 2017; 102:203–16.

39. Castro JP, Wardelmann K, Grune T, Kleinridders A. Mitochondrial chaperones in the brain: safeguarding brain health and metabolism? Frontiers in endocrinology. 2018; 9:196.

40. Ingram T, Chakrabarti L. Proteomic profiling of mitochondria: what does it tell us about the ageing brain? Aging (Albany NY). 2016; 8:3161.

41. Gnaiger E, Arnould T, Detraux D, STORDER J. Mitochondrial respiratory states and rates. MitoFit Preprint Arch (2019). 2019:1–40.

42. Alberts B, Johnson A, Lewis J, Raff M, Roberts K, Wlater P. Molecular Biology of the Cell: Garland Science; 2007.

43. Gnaiger E. Mitochondrial pathways and respiratory control: an introduction to OXPHOS analysis. Bioenergetics communications. 2020; 2020:2-.

44. Liu YJ, McIntyre RL, Janssens GE, Houtkooper RH. Mitochondrial fission and fusion: A dynamic role in aging and potential target for age-related disease. Mechanisms of ageing and development. 2020; 186:111212.

45. Cogliati S, Frezza C, Soriano ME, Varanita T, Quintana-Cabrera R, Corrado M, et al. Mitochondrial cristae shape determines respiratory chain supercomplexes assembly and respiratory efficiency. Cell. 2013; 155:160–71.

46. Zhang W-w, Zhang L, Hou W-k, Xu Y-x, Xu H, Lou F-c, et al. Dynamic expression of glucose transporters 1 and 3 in the brain of diabetic rats with cerebral ischemia reperfusion. Chinese medical journal. 2009; 122:1996–2001.

47. Liu Y, Beyer A, Aebersold R. On the dependency of cellular protein levels on mRNA abundance. Cell. 2016; 165:535–50.

48. Buccitelli C, Selbach M. mRNAs, proteins and the emerging principles of gene expression control. Nature Reviews Genetics. 2020; 21:630–44.

49. Appelman Y, van Rijn BB, Monique E, Boersma E, Peters SA. Sex differences in cardiovascular risk factors and disease prevention. Atherosclerosis. 2015; 241:211–8.

50. Stanhewicz AE, Wenner MM, Stachenfeld NS. Sex differences in endothelial function important to vascular health and overall cardiovascular disease risk across the lifespan. American Journal of Physiology-Heart and Circulatory Physiology. 2018; 315:H1569–H88.

51. Wang S, Zhang H, Liu Y, Li L, Guo Y, Jiao F, et al. Sex differences in the structure and function of rat middle cerebral arteries. American Journal of Physiology-Heart and Circulatory Physiology. 2020; 318:H1219–H32.

52. Gaignard P, Savouroux S, Liere P, Pianos A, Thérond P, Schumacher M, et al. Effect of sex differences on brain mitochondrial function and its suppression by ovariectomy and in aged mice. Endocrinology. 2015; 156:2893–904.

53. Picard M, McEwen BS. Mitochondria impact brain function and cognition. Proceedings of the National Academy of Sciences. 2014; 111:7–8.

54. Hara Y, Yuk F, Puri R, Janssen WG, Rapp PR, Morrison JH. Presynaptic mitochondrial morphology in monkey prefrontal cortex correlates with working memory and is improved with estrogen treatment. Proceedings of the National Academy of Sciences. 2014; 111:486–91.

