## Supplemental Materials for "Differential Left and Right Carotid Artery Blood Flow and Altered Hippocampal Mitochondrial Function After Transverse Aortic Constriction in Aging Rats"

**Supplementary Table 1**: Effects of Condition and Hemisphere on Peak Carotid Blood Flow Velocity.

| | 20 weeks | | | 30 weeks | | | 40 weeks | | |
|---|---|---|---|---|---|---|---|---|---|
| | F (df) | *p* | $\eta p^2$ | F (df) | *p* | $\eta p^2$ | F (df) | *p* | $\eta p^2$ |
| Condition | (1,40)=1.53 | 0.22 | 0.03 | (1,40)=3.95 | 0.054 | 0.09 | (1,37)=9.43 | <.01 | 0.20 |
| Hemisphere | (1,40)=28.1 | <.001 | 0.41 | (1,40)=14.6 | <.001 | 0.26 | (1,37)=19.0 | <.001 | 0.34 |
| Condition*Hemisphere | (1,40)=19.5 | <.001 | 0.32 | (1,40)=15.6 | <.001 | 0.28 | (1,37)=10.8 | <.001 | 0.22 |

**Supplementary Table 2**: Effects of Condition and Hemisphere on Carotid Blood Flow Pulsatility Index.

| | 20 weeks | | | 30 weeks | | | 40 weeks | | |
|---|---|---|---|---|---|---|---|---|---|
| | F (df) | p | $\eta p^2$ | F (df) | p | $\eta p^2$ | F (df) | p | $\eta p^2$ |
| Condition | (1,40)=0.45 | 0.50 | 0.01 | (1,40)=2.59 | 0.11 | 0.06 | (1,37)=7.88 | <.01 | 0.17 |
| Hemisphere | (1,40)=43.1 | <.001 | 0.51 | (1,40)=40.2 | <.001 | 0.50 | (1,37)=75.5 | <.001 | 0.67 |
| Condition*Hemisphere | (1,40)=20.6 | <.001 | 0.34 | (1,40)=22.5 | <.001 | 0.36 | (1,37)=45.9 | <.001 | 0.55 |

**Supplementary Table 3**: Effects of Time on Carotid Artery Hemodynamics and Pulsatility Index.

| | Mean Velocity | | | Peak Velocity | | | Pulsatility Index | | |
|---|---|---|---|---|---|---|---|---|---|
| | F (df) | p | $\eta_p^2$ | F (df) | p | $\eta_p^2$ | F (df) | p | $\eta_p^2$ |
| SHAM-RCCA | (2,27)=0.26 | 0.77 | 0.019 | (2,27)=0.28 | 0.75 | 0.021 | (2,27)=0.08 | 0.91 | 0.007 |
| SHAM-LCCA | (2,27)=0.99 | 0.38 | 0.069 | (2,27)=0.57 | 0.56 | 0.041 | (2,27)=0.26 | 0.76 | 0.019 |
| TAC-RCCA | (2,32)=1.00 | 0.37 | 0.059 | (2,32)=0.79 | 0.45 | 0.048 | (2,32)=0.54 | 0.58 | 0.033 |
| TAC-LCCA | (2,31)=0.20 | 0.81 | 0.013 | (2,31)=0.33 | 0.71 | 0.021 | (2,31)=3.18 | 0.055 | 0.17 |

***Note:*** RCCA- right common carotid artery, LCCA- left common carotid artery, TAC- transverse aortic constriction, SHAM- sham controls.

**Supplementary Table 4**: Effects of Time on Carotid Artery Diameter.

| | Diameter - Systole | | | Diameter - Diastole | | |
|---|---|---|---|---|---|---|
| | F (df) | p | $\eta p^2$ | F (df) | p | $\eta p^2$ |
| SHAM-RCCA | (2,27)=0.40 | 0.67 | 0.029 | (2,27)=0.43 | 0.64 | 0.031 |
| SHAM-LCCA | (2,27)=0.68 | 0.51 | 0.059 | (2,27)=0.35 | 0.70 | 0.026 |
| TAC-RCCA | (2,32)=0.46 | 0.65 | 0.027 | (2,32)=0.46 | 0.63 | 0.028 |
| TAC-LCCA | (2,30)=0.02 | 0.98 | 0.001 | (2,30)=0.06 | 0.93 | 0.005 |

***Note:*** RCCA- right common carotid artery, LCCA- left common carotid artery, TAC- transverse aortic constriction, SHAM- sham controls.

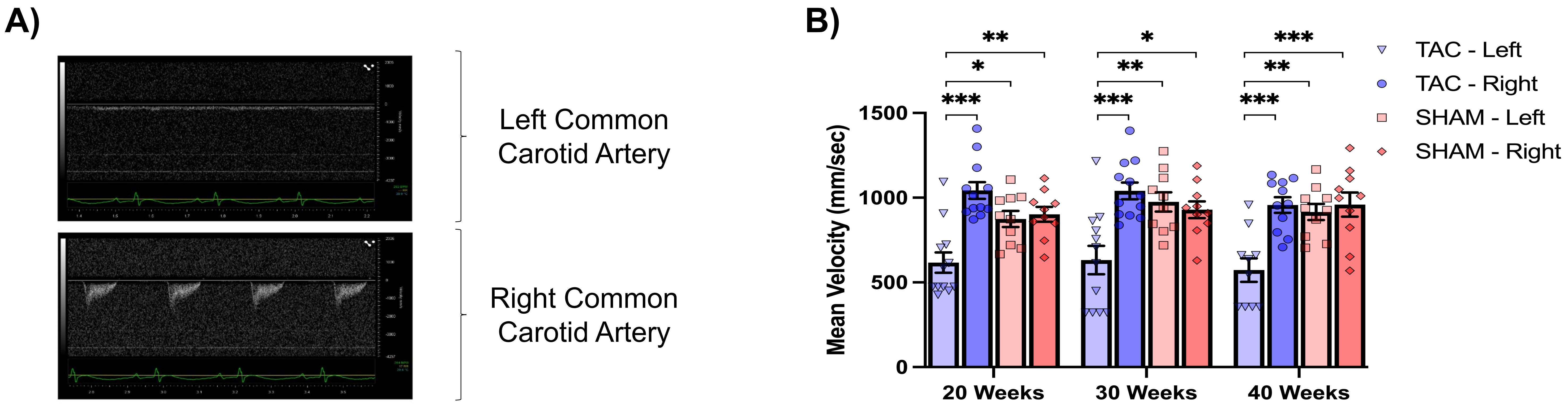


**Supplementary Figure 1**

**Mean Carotid Blood Flow Velocity is Greater in the Right Hemisphere of TAC Animals, and Among Female Animals in TAC and SHAM. A)** Sample exclusionary ultrasound time series displaying severe and unmeasurable left carotid blood flow in TAC animal with measurable carotid blood flow on the right **B)** Mean carotid blood flow velocity in the right and left carotids of TAC and SHAM animals 20-, 30-, and 40 weeks post-surgery. **C)** Sex differences in mean blood flow velocity in TAC animals 20-, 30-, and 40 weeks post-surgery. **D)** Sex differences in mean blood flow velocity in SHAM animals 20-, 30-, and 40 weeks post-surgery. TAC- transverse aortic constriction, SHAM- sham control, mm/sec- millimeter per second, M- male, F- female, * = $p<0.05$, ** = $p<0.01$, *** = $p<0.001$. Data presented as mean + SEM.

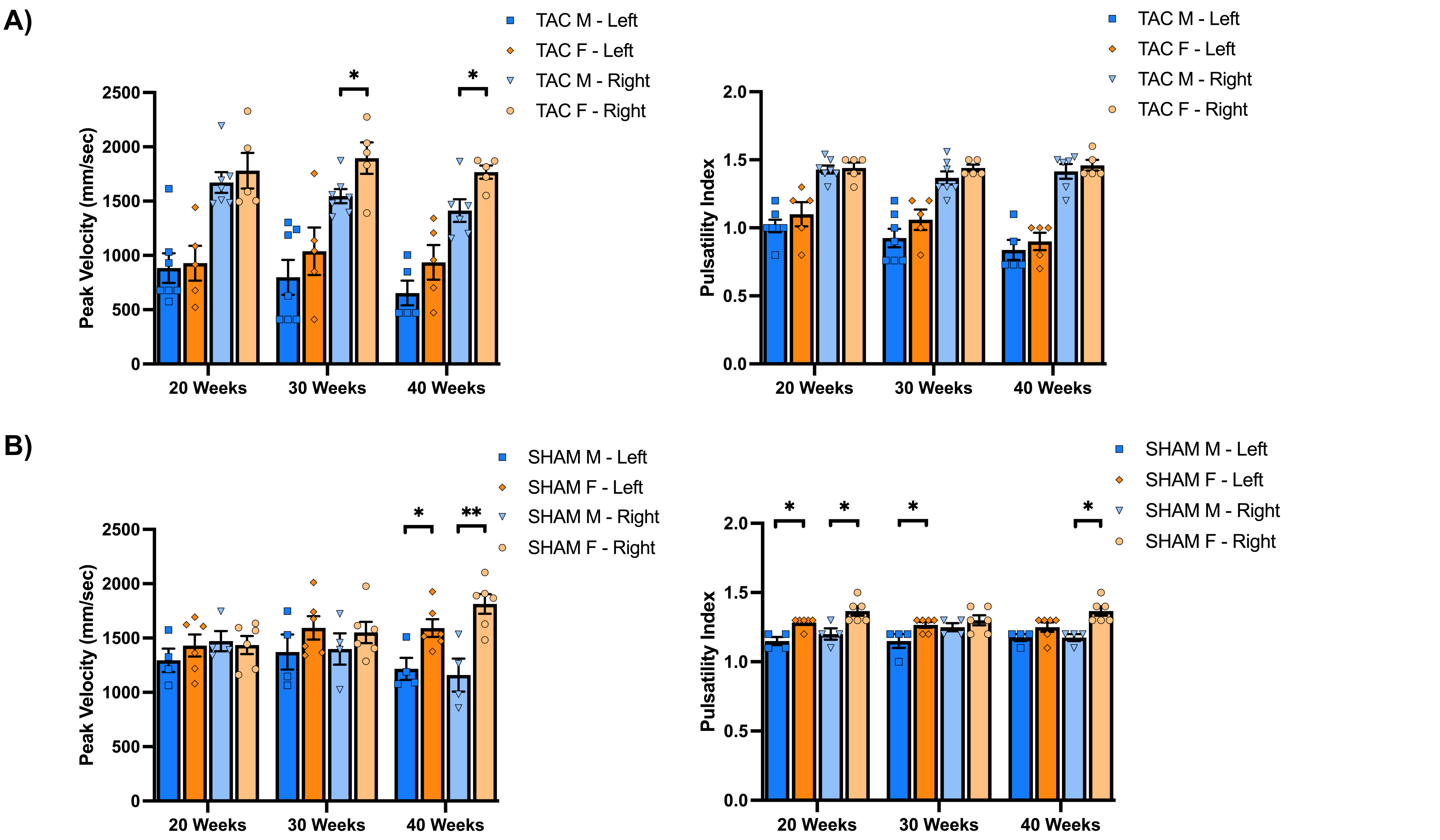


**Supplementary Figure 2**

**Sex Differences in Blood Flow Velocity and Pulsatility are Present in TAC and SHAM Animals. A)** Carotid blood flow velocity and pulsatility in the right and left carotids of male and female TAC animals 20-, 30-, and 40 weeks post-surgery. **B)** Carotid blood flow velocity and pulsatility in the right and left carotids of male and female SHAM animals 20-, 30-, and 40 weeks post-surgery. TAC- transverse aortic constriction, SHAM- sham control, mm/sec- millimeter per second, M-male, F- female, * = $p<0.05$, ** = $p<0.01$. Data presented as mean + SEM.

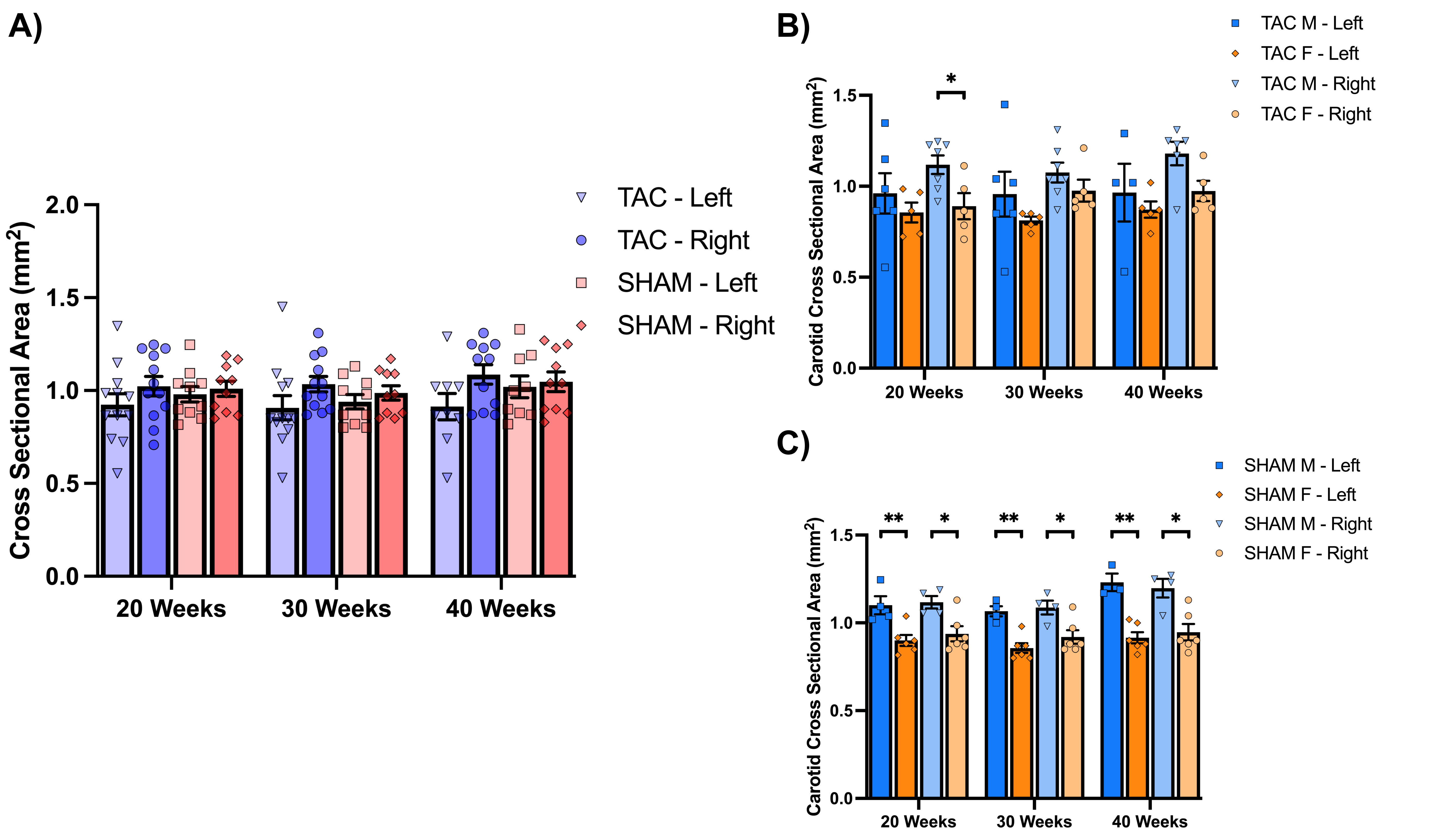


### Supplementary Figure 3

**Carotid Artery Cross-Sectional Does not Significantly Differ Between TAC and SHAM nor by Sex in Either Condition.**
**A)** Calculated carotid cross-sectional area in the right and left carotids of TAC and SHAM animals 20-, 30-, and 40 weeks post-surgery. **B)** Calculated carotid cross-sectional area in the right and left carotids of TAC animals 20-, 30-, and 40 weeks post-surgery. **C)** Calculated carotid cross-sectional area in the right and left carotids of SHAM animals 20-, 30-, and 40 weeks post-surgery. TAC- transverse aortic constriction, SHAM- sham control, $mm^2$- squared millimeters, M- male, F- female. Data presented as mean + SEM.

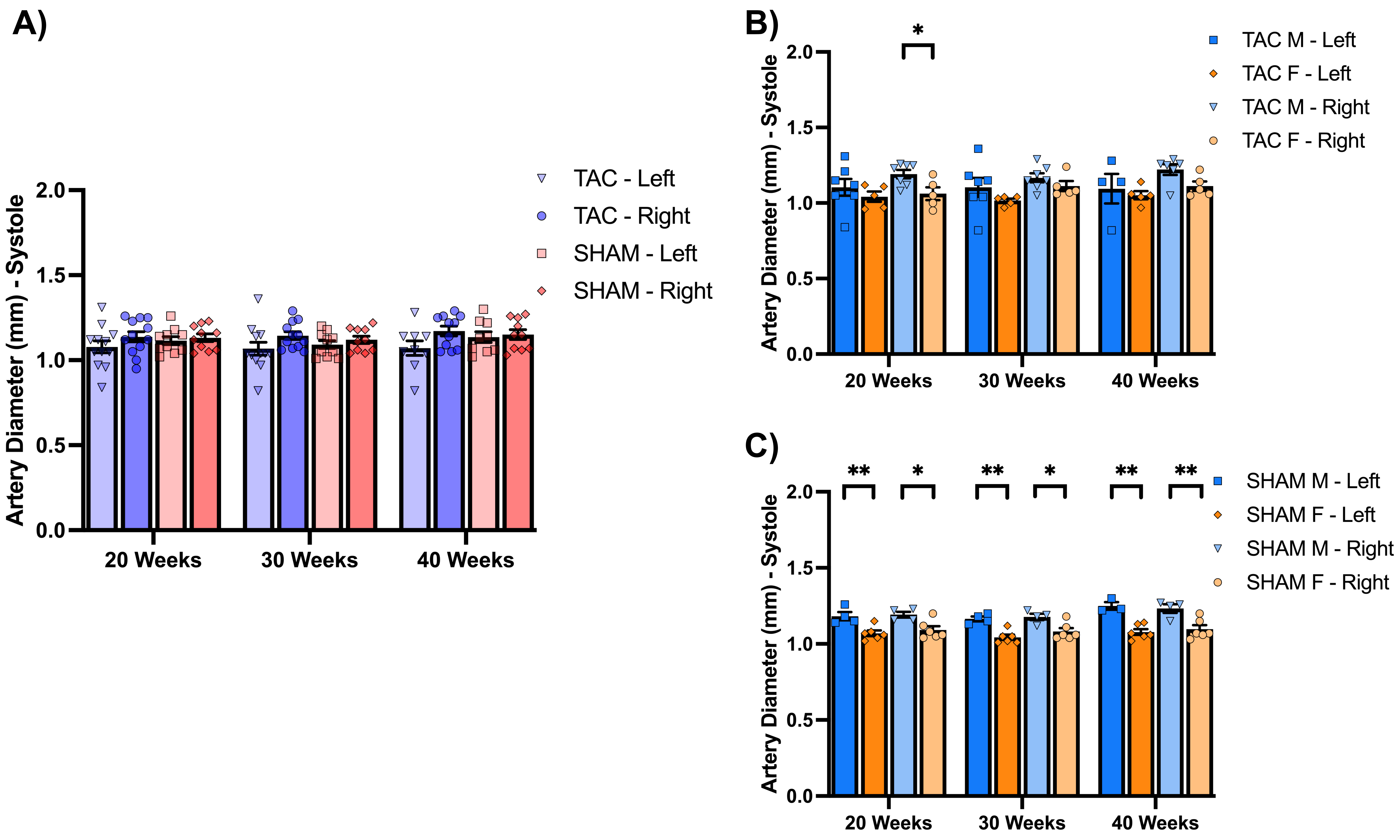


### Supplementary Figure 4

**Sex Differences in Systolic Carotid Artery Diameter are Widespread in SHAM Animals but Constrained to the Right Hemisphere TAC.**
**A)** Systolic carotid artery diameter in TAC and SHAM animals 20-, 30-, and 40 weeks post-surgery. **B)** Systolic carotid artery diameter in male and female TAC animals 20-, 30-, and 40 weeks post-surgery. **C)** Systolic carotid artery diameter in male and female SHAM animals 20-, 30-, and 40 weeks post-surgery. TAC- transverse aortic constriction, SHAM- sham control, mm- millimeters, M- male, F- female, * = p<0.05. Data presented as mean + SEM.

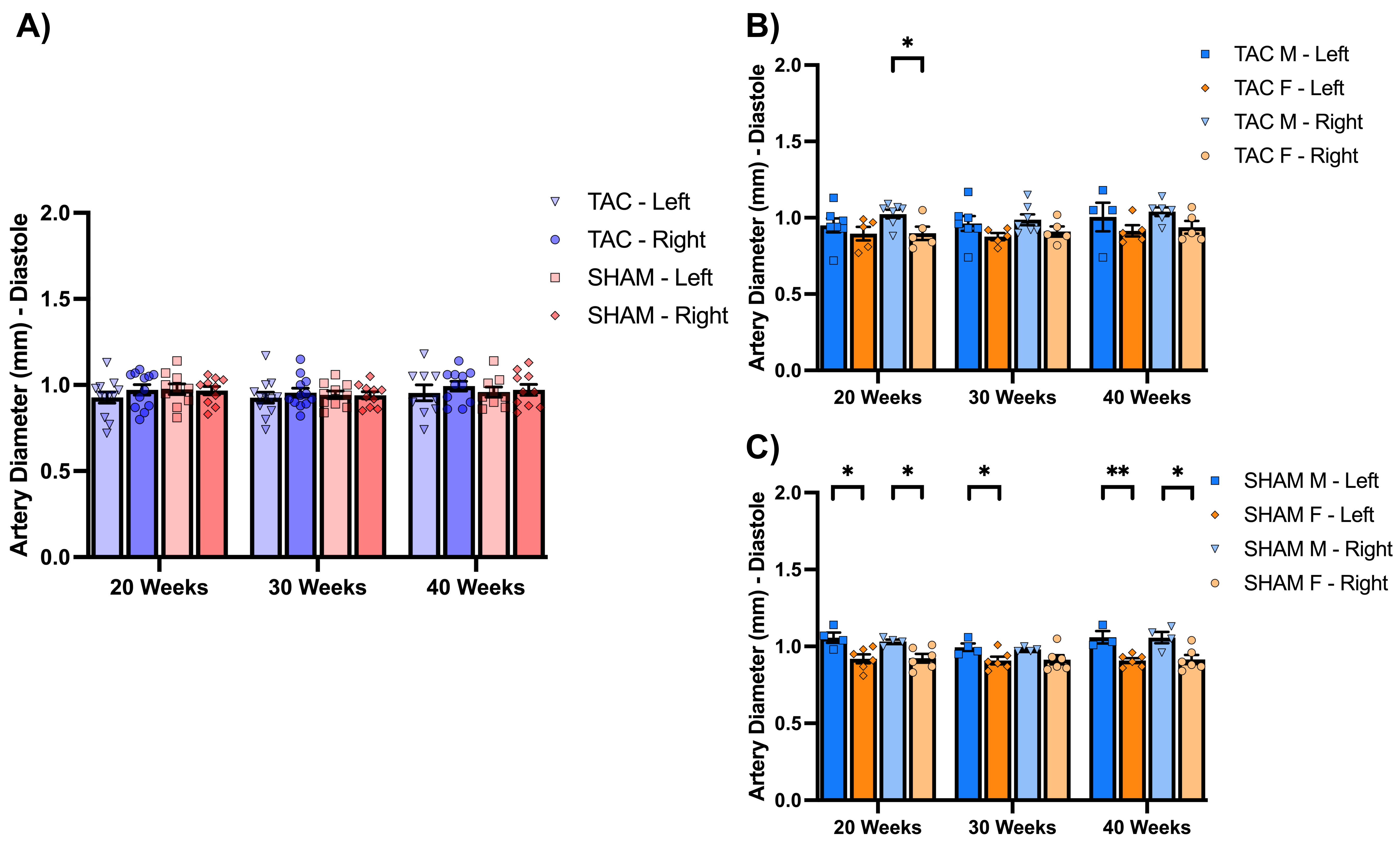


### Supplementary Figure 5

**Sex Differences in Diastolic Carotid Artery Diameter are Widespread in SHAM Animals but Constrained to the Right Hemisphere TAC.**

**A)** Diastolic carotid artery diameter in TAC and SHAM animals 20-, 30-, and 40 weeks post-surgery. **B)** Diastolic carotid artery diameter in male and female TAC animals 20-, 30-, and 40 weeks post-surgery. **C)** Diastolic carotid artery diameter in male and female SHAM animals 20-, 30-, and 40 weeks post-surgery. TAC- transverse aortic constriction, SHAM- sham control, mm- millimeters, M- male, F- female, * = $p<0.05$. Data presented as mean + SEM.

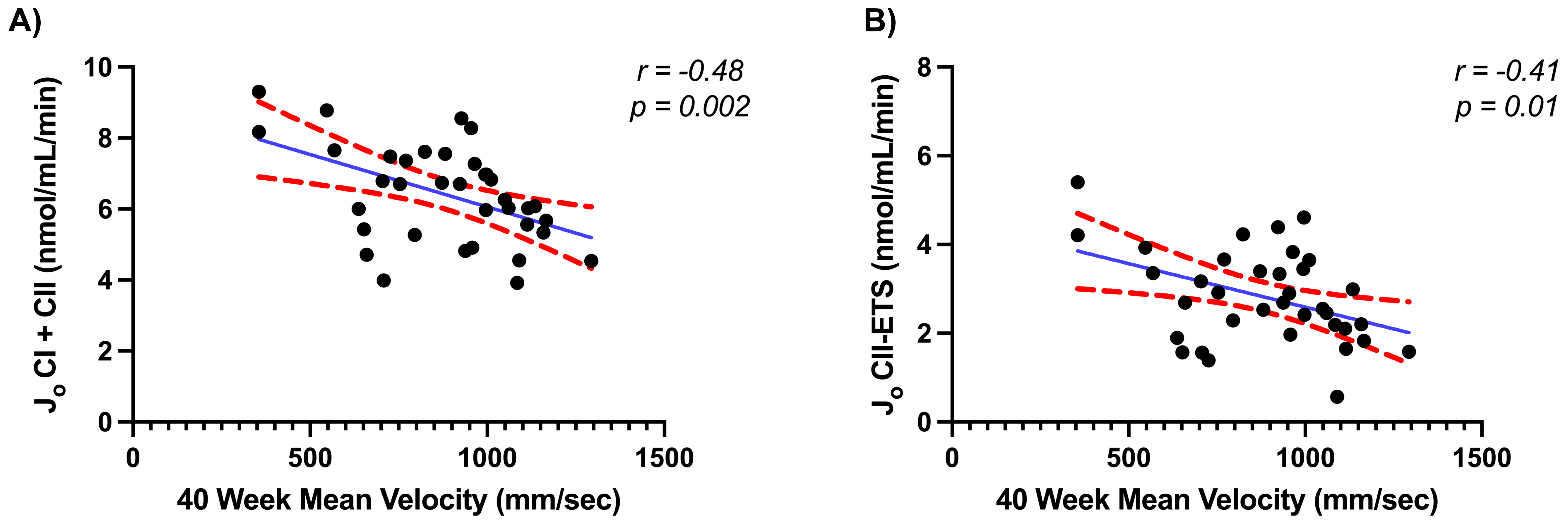


**Supplementary Figure 6**

**High mean carotid blood flow velocity and pulsatility are negatively Associated with Hippocampal Mitochondrial Respiratory States:**
**A)** Scatterplot depicting the negative association between mean carotid artery velocity and CI&II coupled hippocampal respiration. **B)** Scatterplot depicting the negative association between mean carotid artery velocity to CII uncoupled hippocampal respiration. $J_O$ = oxygen consumption, nmol/ml/min = nanomoles per milliliter per minute, CI = complex I, CII = complex II, ETS = electron transfer system.
